# Mild traumatic brain injury alters function in the dorsolateral prefrontal cortex: a TMS-EEG study

**DOI:** 10.1101/2025.05.22.655676

**Authors:** Emily Moore, Emily Webb, Ryoki Sasaki, Nigel C Rogasch, Ngee Foo, John G Semmler, George M Opie

**Affiliations:** Discipline of Physiology, School of Biomedicine, The University of Adelaide, Adelaide, Australia; School of Psychology, Deakin University, Burwood, Victoria, Australia; Hopwood Centre for Neurobiology, Lifelong Health Theme, South Australian Health and Medical Research Institute, Adelaide, Australia; School of Psychology and Turner Institute for Brain and Mental Health, Monash University, Melbourne, Australia; Trauma Service, The Royal Adelaide Hospital, Adelaide, South Australia, Australia

**Author notes:** Correspondence: George M. Opie.

## Abstract

Numerous studies highlight alterations in motor cortical neurophysiology after mild traumatic brain injury (mTBI). In contrast, understanding of effects in cognitive brain areas, which potentially underpin more impactful effects of injury, remains limited. The current study addressed this limitation by using electroencephalography to index responses to transcranial magnetic stimulation (TMS-EEG) applied over the dorsolateral prefrontal cortex (DLPFC). In 18 mTBI patients and 22 healthy controls, TMS-EEG was used to index cortical inhibition and long-term depression (LTD)-like plasticity within DLPFC. These were assessed using paired-pulse TMS (short-[SICI] and long-[LICI] interval intracortical inhibition) and continuous theta burst stimulation (cTBS), respectively. Functional effects of injury were assessed via the N-back and Trail Making Test. Responses were quantified in both temporal (i.e., TMS-evoked potentials; TEPs) and spectral (i.e., TMS-related oscillations; TROs) domains. For TEPs, measures of inhibition and plasticity both tended to be reduced by mTBI. In contrast, TRO data suggested a contradictory increased response of patients to both paired-pulse stimulation and cTBS that was specific to high-frequency bands. In addition, while TMT performance was reduced in the mTBI group, this was unrelated to neurophysiological effects of injury. These findings highlight novel effects of mTBI on DLPFC neurophysiology and emphasise the importance of considering both evoked and induced activity when examining neurophysiological effects of mTBI.

## Introduction

Mild traumatic brain injury (mTBI) can occur following any knock to the head and is often caused by common events like falls, motor vehicle accidents and contact sport. While once considered short-term injuries, we now know mTBI can cause significant and prolonged side-effects in a large proportion of patients. Up to 80% report ongoing symptoms 12 months after injury (McMahon *et al*. 2014), including motor deficits (De Beaumont *et al*. 2009, Pearce *et al*. 2014), increased rates of depression (Kerr *et al*. 2012) and suicide (Fralick *et al*. 2016), a 5-fold increase in the development of mild cognitive impairment and earlier onset of Alzheimer’s disease (Guskiewicz *et al*. 2005). Despite this, we still lack a clear understanding of how the brain is altered after mTBI, which factors determine the clinical trajectory faced by a patient – particularly those that determine rapid resolution vs. prolonged deficit – and what interventions can be applied to influence recovery. Given the high global incidence of these injuries (69 million TBIs annually, ∼80% of which are mTBI; Dewan *et al*. 2018), this demonstrates the pressing need for a better understanding of how mTBI influences the brain.

Although mTBI is rarely associated with structural damage on conventional diagnostic imaging, changes in brain function have been well documented. One method that has been applied frequently in this space is transcranial magnetic stimulation (TMS), a non-invasive brain stimulation technique able to assess excitability within intracortical circuits, and quantify the brain’s ability to change the nature of its synaptic connectivity (i.e., neuroplasticity; Suppa *et al*. 2022). Conventional application of TMS involves stimulation over primary motor cortex (M1), with responses recorded in peripheral muscles providing indirect measures of cortical excitability. Using this approach, several studies have demonstrated injury-related changes in intracortical inhibitory circuits involving gamma aminobutyric acid (GABA) receptors (De Beaumont *et al*. 2009, Tremblay *et al*. 2011, De Beaumont *et al*. 2012, Miller *et al*. 2014, Pearce *et al*. 2014, Di Virgilio *et al*. 2016, Edwards and Christie 2017) receptors, as well as an altered neuroplastic response (De Beaumont *et al*. 2012, Pearce *et al*. 2018). These changes have been observed across broad post-injury timeframes ranging from days (Pearce *et al*. 2015) to decades (De Beaumont *et al*. 2009, Pearce *et al*. 2014), and have been associated with deficits in motor learning and performance (De Beaumont *et al*. 2012).

While conventional TMS has provided important neurophysiological insights into mTBI, the direct relevance of this information is limited to the specific cortical circuits that contribute to generation of the MEP. In contrast, effects of mTBI involve brain networks that extend beyond M1 (Morelli *et al*. 2021), and understanding the neurophysiological changes within these areas is likely to provide important insight into the cognitive effects of injury. One approach to address this limitation is to use electroencephalography to record the response to TMS (TMS-EEG) when applied over non-motor brain areas. This generates a TMS-evoked EEG potential (TEP), the amplitude and spatio-temporal characteristics of which reflect activity of cortical circuits within the stimulated cortex and interconnected areas (Hernandez-Pavon *et al*. 2023). Several recent studies have used this approach to investigate effects of mTBI within the dorsolateral pre-frontal cortex (DLPFC). Interestingly, results suggested an injury-related decrease in the excitation-inhibition balance (Coyle *et al*. 2023b, Coyle *et al*. 2024), in addition to changes in the neuroplastic response (Coyle *et al*. 2023a). Furthermore, the nature of these effects varied over the post-injury period (Coyle *et al*. 2023b, Coyle *et al*. 2023a).

These studies demonstrate the utility of TMS-EEG for quantifying neurophysiological effects of mTBI in DLPFC. In addition, they also suggest that further examination of injury-related changes in this area is warranted. The current study therefore sought to build on the existing work in two ways. First, it is possible to use TMS-EEG to record the response to the paired-pulse TMS measures of short-(SICI) and long (LICI) interval intracortical inhibition. While effects of these paradigms on the TEP are unlikely to directly reflect the mechanisms that have been defined in motor cortex with conventional TMS (Premoli *et al*. 2018), they nonetheless provide an additional index of neuronal activation/modulation that might provide unique insights to the effects of mTBI. Second, previous assessment of neuroplasticity was limited to an intervention thought to induce long-term potentiation (LTP)-like effects, whereas previous work from our group (Opie *et al*. 2019) and others (De Beaumont *et al*. 2012) suggests mTBI also influences long-term-depression (LTD)-like mechanisms. Consequently, the current study used TMS-EEG applied to DLPFC to assess mTBI-related changes in the response to SICI and LICI, and continuous theta burst stimulation (cTBS; thought to induce LTD-like changes). To investigate how neurophysiological effects of injury relate to functional deficits, performance on the N-back and Trail Making Test was also related to TMS-EEG measures; these tests involve DLPFC (Owen *et al*. 2005, Zakzanis *et al*. 2005) and are sensitive to mTBI (Arciniega *et al*. 2021, Li *et al*. 2023).

## Methods

### Participants

A total of 40 participants were recruited, including 18 that had sustained an mTBI within the previous 24 months (mean age ± SD: 33.1 ± 13.2 years) and 22 healthy controls (28.8 ± 8.19) with no history of head injury. mTBI was defined as a head injury causing any of: *(1)* a Glasgow Coma Scale (GCS) score of 13-15 30 minutes after injury; *(2)* a period of post-traumatic amnesia not greater than 24 hours or *(3)* a loss of consciousness no greater than 30 minutes. Recruitment of mTBI patients was through the Royal Adelaide Hospital Trauma Service, and both patients and healthy controls were recruited from within the University and wider community, through word of mouth, flyers, and social media advertising.

All participants were screened using an established questionnaire (Rossi *et al*. 2011), with exclusion criteria including: history of brain lesion; history of neurological disease; history of alcohol or substance abuse, or psychiatric illness; cardiac pacemakers; metal implants within the skull; pregnancy; medications affecting cortical excitability or history of more severe brain injury. The study was approved by both the Central Adelaide Local Health Network and the University of Adelaide Human Research Ethics Committees (approval HREC/17/RAH/116). Each participant provided written, informed consent prior to participation.

### Experimental Arrangement & Recordings

Each experimental session was separated by at least 1 week, and the order of sessions was randomised between participants. At the start of each session, current symptomatology was assessed using a standardised post-concussion symptom (PCS) questionnaire^11^ (rating current experience of 21 symptoms from none [0] to severe [6]), whereas tests of executive function were completed via a computerised platform (detailed below). For the remainder of the session, participants were then seated comfortably, with hands supported on a cushion in their lap.

*Electromyography (EMG):* As intensity of DLPFC stimulation was titrated against motor cortical thresholds, EMG was used to record motor evoked potentials in response to stimulation of M1. Two Ag-AgCl electrodes were placed on the skin over the first dorsal interosseous (FDI) muscle in a belly-tendon montage, with a third electrode placed over the Ulna styloid process grounding the signal. EMG was amplified (1000x) and band-pass filtered (20Hz-1kHz) using a CED1902 (Cambridge Electronic Design, Cambridge, UK) prior to digitisation at 2kHz through a CED1401 interface (Cambridge Electronic Design, Cambridge, UK). Signal noise associated with mains power was reduced using a Humbug mains power noise eliminator (Quest Scientific, North Vancouver, Canada).

*Electroencephalography (EEG):* EEG data was recorded using a WaveGuard EEG Cap (ANT Neuro, Hengelo, The Netherlands), with 62 sintered Ag/Ag-Cl electrodes in standard 10-10 positions, connected to an eego mylab amplifier (ANT Neuro). The AFz electrode was used to ground the recordings, whereas CPz was used as reference for all electrodes. EEG signals were sampled at 8kHz and filtered online (DC-0.26 x sampling frequency). The electrode impedance was maintained at less than 10 kΩ for the duration of all recordings. During collection of EEG, participants were instructed to keep their face relaxed, and to avoid blinking where possible. Auditory input during stimulation was minimised by playing white noise through in-ear headphones, with volume adjusted until TMS could no longer be perceived, or the upper limit of comfort was reached.

### Tests of Executive function

Participants completed two computerised tests (via the Psychology Experimental Building [PEBL] platform; Mueller and Piper 2014) of executive function at the beginning of each session, including:

**-Trail Making Test (TMT):** In a series of 2 subtests, 26 consecutive numbers (TMT-A), or 13 consecutive numbers and 13 consecutive letters (TMT-B), were displayed across the screen. For each test, participants were required to click on each item in sequence as quickly and accurately as possible. For TMT-B, this required alternating selection of numbers and letters (i.e., 1 – A – 2 – B – …). Performance within each subtest was assessed via the mean time to completion (seconds) and accuracy (defined as the number of clicks on screen *vs.* number of targets).
**-N-Back**: A sequence of consonants was presented successively at the centre of a screen. Participants were required to respond via a keyboard press when the displayed letter was the same as the one shown *N* letters prior. Difficulty levels of *N* = 1 and 2 were assessed, with 102 trials collected for each condition, a target rate of 25% and interstimulus interval of 2s. Performance was assessed using reaction time for correct trials, in addition to the *d*-prime metric (i.e., the *Z*-transformed difference between the rate of hits and false-alarms)(Haatveit *et al*. 2010).

### Transcranial Magnetic Stimulation

For both sessions, all TEP data were collected using a branding iron figure-of-eight coil connected to two Magstim 200^2^ units via a Bistim module (Magstim, Dyfed, UK). The coil was held tangentially to the scalp and at an angle of 45 degrees to the sagittal plane, resulting in an anteriorly directed current flow in the brain. Stimulation was initially applied over the hand representation of M1, optimised to generate the largest and most consistent response within the right FDI following moderately suprathreshold TMS. Resting motor threshold (RMT) was identified for this location, corresponding to the stimulus intensity (% maximum stimulator output; %MSO) producing an MEP amplitude ≥ 50 µV in at least 5 out of 10 consecutive trials. Following identification of rMT, all subsequent stimulation was applied over the left DLPFC, identified as the midpoint between EEG electrodes F3 and F5 (Rusjan *et al*. 2010). For both M1 and DLPFC sites, location of stimulation was logged relative to the MNI-ICBM152 head model using neuro-navigation software (Brainsight 2, Rogue, Canada). To avoid anticipation of the stimulus, all non-interventional TMS was applied at a rate of 0.2 Hz ± 10%. All TMS was applied with the EEG cap in place.

*SICI & LICI:* SICI was assessed using a 70% rMT conditioning stimulus, 120% rMT test stimulus and interstimulus interval (ISI) of 2 ms. LICI was assessed with conditioning and test stimuli set at 120% rMT and ISI of 100 ms. Single-pulse stimuli were additionally applied at 120% rMT and 70% rMT: while the former allowed quantification of cortical excitability, the latter was used for correction of the TEP generated by SICIs conditioning stimulus (see below). As 104 stimuli were applied for each condition, assessment of ICI involved a total of 416 stimuli. To avoid a loss of participant attention, this was applied over 8 blocks of 52 stimuli, with each block containing equal numbers of each stimulus condition applied in a pseudo-randomised order.

*Continuous theta burst stimulation:* Continuous TBS involved bursts of three stimuli applied at 50 Hz that were repeated at 5 Hz for 40s, totalling 600 pulses^32^. This was applied over DLPFC at an intensity of 70% rMT using a Super Rapid stimulator (Magstim, Dyfed, UK). To avoid loss of stimuli, intensity was capped at 52% MSO for all participants, resulting in capped intensities for 7 mTBI patients and 13 healthy controls. Stimulus intensity was recalibrated over M1 for this machine, due to the different coil geometry and stimulus waveform (biphasic). Effects of cTBS within DLPFC were quantified by recording single-pulse TEPs before (Pre), directly after (Post 0) and 30 mins after (Post 30) application of the intervention. These data were collected using the branding iron coil and Magstim 200^2^ stimulators (see above). Each TEP block was applied at 120% rMT, involved 104 stimuli and was recorded as two blocks of 52 to avoid loss of attention.

### Data Analysis

EEG data was analysed using MATLAB (R2021b, The Mathworks, USA) with EEGLAB (Delorme and Makeig 2004), Fieldtrip (Oostenveld *et al*. 2010) and the TMS-EEG signal analyser (TESA) (Rogasch *et al*. 2017) toolboxes, whereas figures were generated using the matplotlib toolbox in python (v3.11).

*Pre-processing:* For recordings from both sessions, data were epoched around the TMS pulse (±1500ms) and baseline corrected (cTBS session: −1000 to −10 ms; ICI session: −1000 to −110 ms). The peak of the artifact produced by TMS was then removed by cutting the data from −6 to 13 ms (corresponding to the test stimulus in all conditions and encompassing SICIs conditioning and test stimulus), and −110 to −50 ms (LICI condition only, removing the conditioning stimulus artefact) and replacing the excised section using cubic interpolation.

Data were consequently downsampled to 1000 Hz and visually inspected for noisy electrodes and epochs. Given the differences in TMS artefact for each of the ICI conditions, data from each stimulus paradigm were split into separate blocks for the remainder of the analysis. In contrast, data from the TBS session remained concatenated. The source utilised noise-discarding (SOUND) filter was then applied to reduce the large artifacts associated with TMS, in addition to replacing missing electrodes (Mutanen *et al*. 2018, Mutanen *et al*. 2020). A regularisation parameter of 0.1 was used and 10 iterations were applied. An initial round of independent component analysis (ICA) was then run using the FastICA algorithm (Hyvärinen and Oja 2000) and independent components (ICs) representing the tail of the TMS artefact were removed. High-pass (> 1 Hz) and notch (48-52 Hz) butterworth filters were then applied using the *tesa_filtbutter* function (zero phase, fourth order), prior to application of a second round of Fast ICA. ICs associated with ocular artifacts and persistent EMG activity were identified using the *tesa_compselect* function and removed after manual confirmation. Data were then band-pass filtered with *tesa_filtbutter* (1-100 Hz), baseline corrected (−1000 to −10 ms) and re-epoched (±1000 ms) to remove any boundary effects.

For ICI data, both conditioning and test stimuli generate TEPs, the interaction of which can obscure the response to paired-pulse stimulation. Consistent with previously described methods (Opie *et al*. 2018, Opie *et al*. 2019, Sasaki *et al*. 2023), we therefore implemented a subtraction procedure to remove the TEP generated by the conditioning stimulus. This was achieved by aligning the single pulse TEPs applied at 120% rMT (for LICI) or 70% rMT (for SICI) to coincide with the timing of each conditioning stimulus (-100 ms for LICI; −2 ms for SICI) and subtracting them from the paired-pulse data. Given the limitations of this linear approach to potentially non-linear interactions, results for non-corrected data are additionally reported in supplementary materials.

*TEP quantification:* TEP data were quantified locally, using a region-of-interest (ROI) analysis involving electrodes beneath the coil (i.e., averaged over F3 and F5 electrodes), as well as across all electrodes (global analysis). For both local and global analyses, group-averaged data from baseline (TBS session) or test alone (ICI session) conditions was inspected to identify timing of commonly investigated TEP peaks, revealing early (P30, N45, P60) and late (N100, P180) components. However, given the high likelihood of sensory contributions to the late components (Biabani *et al*. 2019, Conde *et al*. 2019, Biabani *et al*. 2021, Gordon *et al*. 2021, Rocchi *et al*. 2021, Biabani *et al*. 2024), TEP quantification focussed exclusively on the early peaks. As the timing of the early components was consistent across sessions and groups, quantification of TEP amplitude used consistent time periods across all stimulus conditions. For ROI analyses, the timing of each peak was first identified in group averaged test alone/Pre data as the maximum positive peaks between 22 – 37 ms (P30) and 53 – 67 ms (P60) and maximum negative peak between 38 – 52 ms (N45). The amplitude of each peak was then calculated as the average of the signal ± 5 ms relative to the timing of the peak, corresponding to 28 – 38 ms (P30), 39 – 49 ms (N45) and 56 – 66 ms (P60). These periods were then used for quantification of peak amplitudes within paired-pulse and post cTBS data. For global analyses, component amplitudes were examined across slightly broader time windows (P30: 22 – 37 ms; N45: 38 – 52 ms; P60: 53 – 67 ms) that better encapsulated peak timing across the scalp. Individual changes in TEP amplitude due to paired-pulse stimulation or cTBS were quantified relative to the size of the test TEP or baseline TEP, respectively. This was calculated using the approach applied in our previous work (Opie *et al*. 2018, Opie *et al*. 2019), expressing the difference in TEP amplitude between conditions (single-pulse/baseline *vs.* paired-pulse/post TBS) as a percentage of the single-pulse/baseline TEP peak-to-peak amplitude in the 13 to 220 ms period.

*TMS-related oscillatory activity (TROs)*: Spectral perturbation associated with TMS in each condition was quantified using time-frequency decomposition. This was achieved with a multitaper approach, using the *mtmconvol* option of Fieldtrip’s *ft_freqanalysis* function. A Hanning taper with frequency-dependent window length was applied (width: 3 cycles per window; time step: 1 ms; frequency step: 1 Hz) to individual trial data for each electrode, producing an estimate of total power (Roach and Mathalon 2008). For all single-pulse conditions, power values were then baseline corrected relative to the −800 to −400 ms pre-stimulus period. However, for paired-pulse conditions, the use of individual trial data meant that an influence of the conditioning stimulus remained present in our estimates of total power. Prior to baseline correction, the paired-pulse correction procedure was therefore repeated for SICI and LICI in the frequency domain. Following this, normalised indices of changes in oscillatory power associated with paired-pulse stimulation, or application of cTBS, were quantified. This was achieved by quantifying the percentage change in the area under the power curve (AUC) for paired-pulse and post-TBS data, relative to single-pulse and pre-TBS data.

### Statistical Analysis

*TEP data – ROI analysis:* Raw TEP data for each peak (P30, N45, P60) were compared between stimulus conditions (ICI: Test, SICI, LICI; cTBS: Pre, Post 0, Post 30) and groups (HC, mTBI) using Bayesian generalised linear mixed models (GLMM). For analysis of spectral data, TRO power was averaged over frequency bands (theta: 5-7 Hz; alpha: 8-13 Hz; lower (l) beta: 14-20 Hz; upper (u) beta: 21-30 Hz; gamma: 31-45 Hz) and time (14-220 ms for frequencies < 30 Hz; 14-120 ms for frequencies > 30 Hz). Values were then compared between stimulus conditions and groups using Bayesian GLMMs. While most ROI analyses were modelled using a skewed normal distribution with identity link function, effects of paired-pulse stimulation on normalised alpha power were modelled with a gaussian distribution and identity link function.

*TEP data – Global analysis:* TEP data for each peak (P30, N45, P60) were compared between stimulus conditions (ICI: Test, SICI, LICI; cTBS: Pre, Post 0, Post 30) and groups (HC, mTBI) using cluster-based permutation analyses. For between-group comparisons of the response to single-pulse stimulation, data from individual subjects was first pooled across sessions, given consistent stimulation parameters used in each (i.e., both Test and Pre were applied at 120% RMT). Clusters were defined as 2 or more neighbouring electrodes having *t*-values with an associated *P*-value < 0.05. Identified clusters were then compared to a permutation distribution generated using the Monte-Carlo method (10,000 permutations) and deemed significant if their cluster-statistic (sum of all t-values in the cluster) exceeded *P* < 0.05 (Maris and Oostenveld 2007). Temporal data were averaged over the target period for each peak of interest (defined above), whereas spectral data were averaged over frequency bands and time as defined for ROI analyses.

To examine the relationship between cortical function and effects of injury, Spearman’s rank correlations were used to correlate normalised TEP data (both temporal and spectral domains) with scores from *N-back* and *TMT*, in addition to time-since-injury. This was performed at all electrodes using a cluster-based permutation approach consistent with our previous work (Sasaki *et al*. 2023). Given the large number of comparisons made, we implemented additional control against false positives by applying a false-discovery rate correction (Benjamini and Hochberg 1995, Martinez-Cagigal 2025) within each stimulus condition (e.g., LICI, SICI, Post 0, Post 30), separately for temporal and spectral data.

*Executive function data:* Performance on *N*-back (indexed via reaction time and *d*‘ scores for each *N*-level) and TMT (indexed via completion time and accuracy for each sub-test) was compared between groups (HC, mTBI) and sessions (ICI, TBS) using Bayesian GLMMs. All models included session order as a factor. A gamma distribution with log link function was used to model reaction time data for both *N*-back and TMT, in addition to d‘ values derived from *N*-back. For TMT accuracy data, a zero-one-inflated beta distribution with logit link function was applied to address data clustering around a value of 1. Due to technical issues during data collection, some participants had missing data (3.8% across all tests). To compensate for this, multiple imputation was applied using the *mice* package (Van Buuren and Groothuis-Oudshoorn 2011), with 20 imputed datasets generated.

*Bayesian estimation:* All Bayesian GLMMs were computed in R (v 4.4.3) using R-studio (v2024.12.1). Posterior distributions for all models were estimated using the No-U-Turn-Sampler (NUTS) extension of Hamiltonian MCMC, implemented within the BRMS package (Bürkner 2017). Each model was run using 8 independent chains, with 1000 warm up and 6000 post-warm up samples (totalling 48,000 post warm-up samples), and default flat priors. Chain convergence was assessed using Rhat values < 1.1 and posterior predictive checks (Gelman and Rubin 1992, Gabry *et al*. 2019). All models implemented the maximal random effect’s structure allowed by the data (i.e., by-participant random intercepts and slopes) in the first instance (Barr 2013). Any model demonstrating issues with chain convergence was re-run with additional iterations (increased to 20,000 per chain). For models failing to converge after this point, the random effect’s structure was simplified to by-participant random slopes.

Custom contrasts of main effects and interactions were generated using the *emmeans* package (Lenth 2025), with effect existence and significance idenitified using the probability of direction (pd; Makowski *et al*. 2019) and region of practical equivalence, respectively. For the current study, ROPE was defined as ± 5% of the standard deviation (on the link scale) for the data being compared (Kruschke 2018). The null hypothesis of no difference was accepted for comparisons where the 89% highest density interval (HDI) fell completely inside the ROPE. In contrast, the null hypothesis was rejected when the 89% HDI fell completely outside the ROPE. No decision was made if the 89% HDI partially overlapped ROPE (Kruschke 2018, Puri *et al*. 2023, Opie *et al*. 2024). Results for all Bayesian models are presented as median values [lower 89% HDI, upper 89% HDI].

## Results

Due to technical difficulties, data from one mTBI patient was not able to be used for analysis. Furthermore, an additional mTBI patient and two healthy controls only attend a single session. Consequently, the final cohort consisted of 37 participants, with baseline characteristics for this group shown in Table 1. While PCS was significantly higher in the mTBI group (*P* = 0.01), no differences were found for all other variables (all *P*-values > 0.05). In addition, no differences in the TEP produced by single-pulse TMS were found between groups, and this was consistent across all comparisons in both temporal and spectral domains.

**Table 1.** Participant characteristics.

| **Table 1. Participant characteristics** |  |  |
| --- | --- | --- |
|  | **HC** | **mTBI** |
| **Sample (n)** |  |  |
| *ICI* | 20 | 17 |
| *cTBS* | 21 | 16 |
| **Age (Years)** |  |  |
| *ICI* | 28.4 ± 8.2 | 31.2 ± 12.3 |
| *cTBS* | 28.1 ± 8.2 | 31.6 ± 12.0 |
| **RMT (% MSO)** |  |  |
| *ICI* | 64.4 ± 10.2 | 61.3 ± 15.2 |
| *cTBS* | 64.3 ± 8.7 | 60.7 ± 12.0 |
| **Days since injury** |  |  |
| *Mean* | NA | 175.3 ± 139.3 |
| *Range* | NA | 9 – 507 |
| **Previous mTBI** |  |  |
| *Mean* | NA | 1.8 ± 1.7 |
| *Range* | NA | 1 – 7 |
| **PCS** | 7.3 ± 9.1 | 20.9 ± 21.9* |

### SICI and LICI

#### ROI analysis

Figure 1 shows results of ROI analyses from the ICI session, including data from both temporal (Fig 1A/1B) and spectral (Fig 1C/1D) domains. Note that the results reported below refer to data that has undergone paired-pulse correction, whereas uncorrected data are reported in supplementary results. For P30 in healthy controls, SICI was associated with a consistent (*pd* = 96.6%) and significant (0% in ROPE) reduction in amplitude, relative to test alone. In addition, LICI produced consistent (*pd* = 96.3%) increases in amplitude, relative to test, but these failed to reach a practical level of significance (0.2% in ROPE). For N45 in healthy controls, LICI was associated with a reduction in amplitude (i.e., less negative) which was consistent (*pd* = 97.4%) and significant (0% in ROPE). Furthermore, comparisons of normalised data showed a relatively consistent (*pd* = 91%) decrease in LICI of N45 when comparing mTBI patients to controls, but this failed to reach a practical level of significance (5% in ROPE). All other comparisons in the time domain were inconsistent and failed to provide sufficient evidence to reject or accept the null hypothesis (all *pd* < 90.2%, % in ROPE: 3.3 – 7.7).

**Figure 1.**
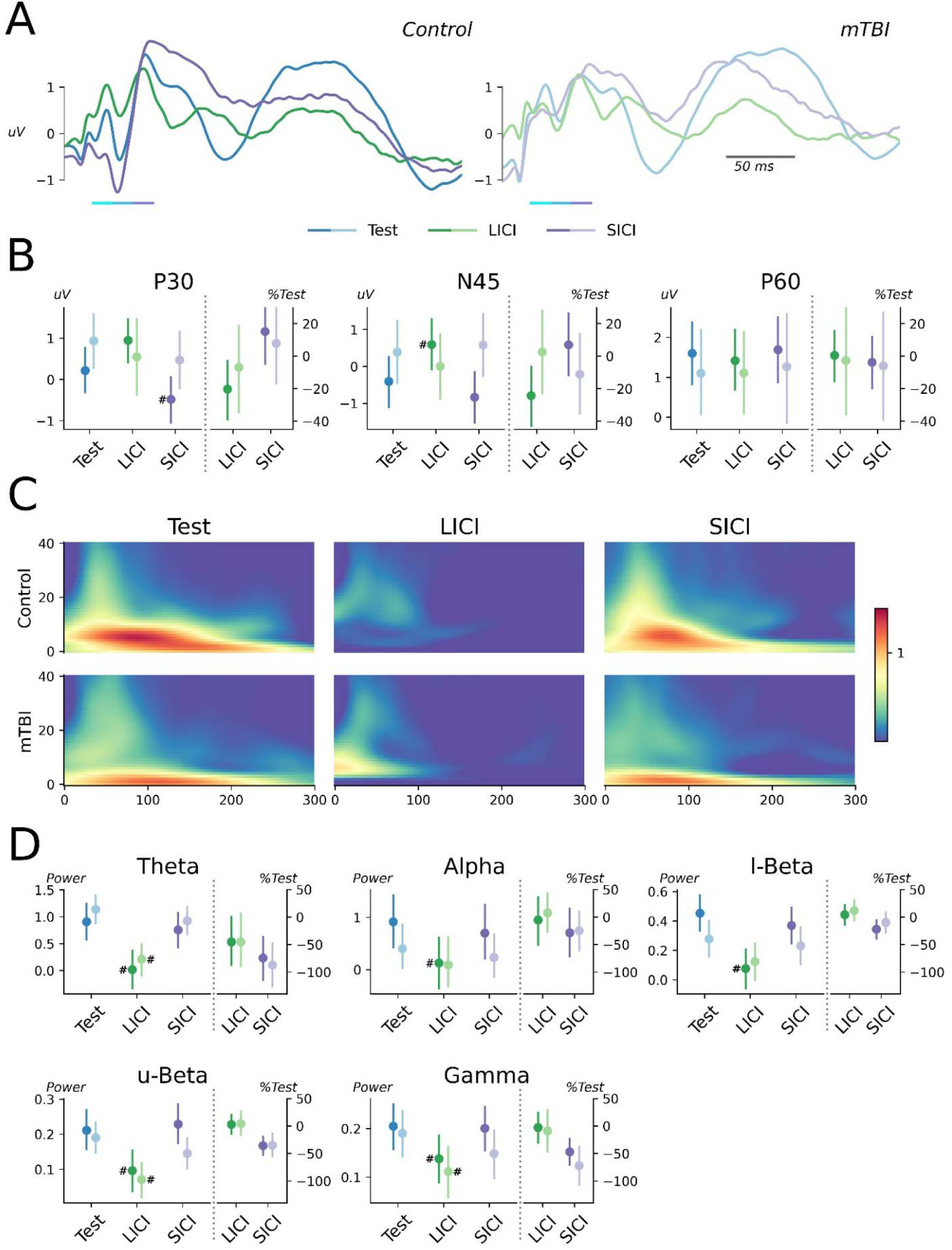
Results of ROI analysis from ICI session. **(A)** TEPs from F3-F5 electrodes recorded in response to test alone *(blue line)*, LICI *(green line)* and SICI *(purple line)* in healthy controls *(left panel)* and mTBI patients *(right panel)*. Coloured bars below line plots highlight time periods used for quantification of TEP peaks. **(B)** Amplitude of P30 *(left panel)*, N45 *(middle panel)* and P60 *(right panel)* for each stimulus condition, compared between healthy controls *(dark colours)* and mTBI patients *(pale colours)*. Raw values are compared on the left y-axis, normalised values are compared on the right y-axis **(C)** Time-frequency responses generated by application of test alone *(left panels)*, LICI *(middle panels)* and SICI *(right panels)* for healthy controls *(top row)* and mTBI patients *(bottom row)*. **(D)** TRO power in theta, alpha, lower beta *(top row)*, upper beta and gamma *(bottom row)* bands for each stimulus condition, compared between healthy controls and mTBI patients. Raw values are compared on the left y-axis, normalised values are compared on the right y-axis. ^#^Significant and consistent difference relative to test alone.

For TROs, application of LICI was associated with reduced power relative to test alone in theta, upper beta and gamma bands for both groups and this was consistent (all *pd >* 97.4%) and significant (0% in ROPE for all comparisons). LICI in healthy controls was also associated with consistent and significant reductions in power for alpha and lower beta bands (all *pd* > 99.6%, 0% in ROPE for all comparisons). All other comparisons in the spectral domain were inconsistent and failed to provide sufficient evidence to reject or accept the null hypothesis (all *pd* < 92.1%, % in ROPE: 5.6 – 24.4).

#### Global analysis

Figure 2 shows grand average TEP waveforms and associated scalp topographies for each stimulus condition from the ICI session, separately for healthy control (Fig 2A) and mTBI patients (Fig 2B); results of within-subject comparisons between single- and paired-pulse data are shown in figure 2C/D. Relative to the Test response, application of SICI in the control group increased P30 (negative cluster: *P* = 0.04; Fig 2C). No other differences were found between single- and paired-pulse responses for either group. Furthermore, comparisons of the normalised response to paired-pulse stimulation failed to identify any significant clusters (Fig 3).

**Figure 2.**
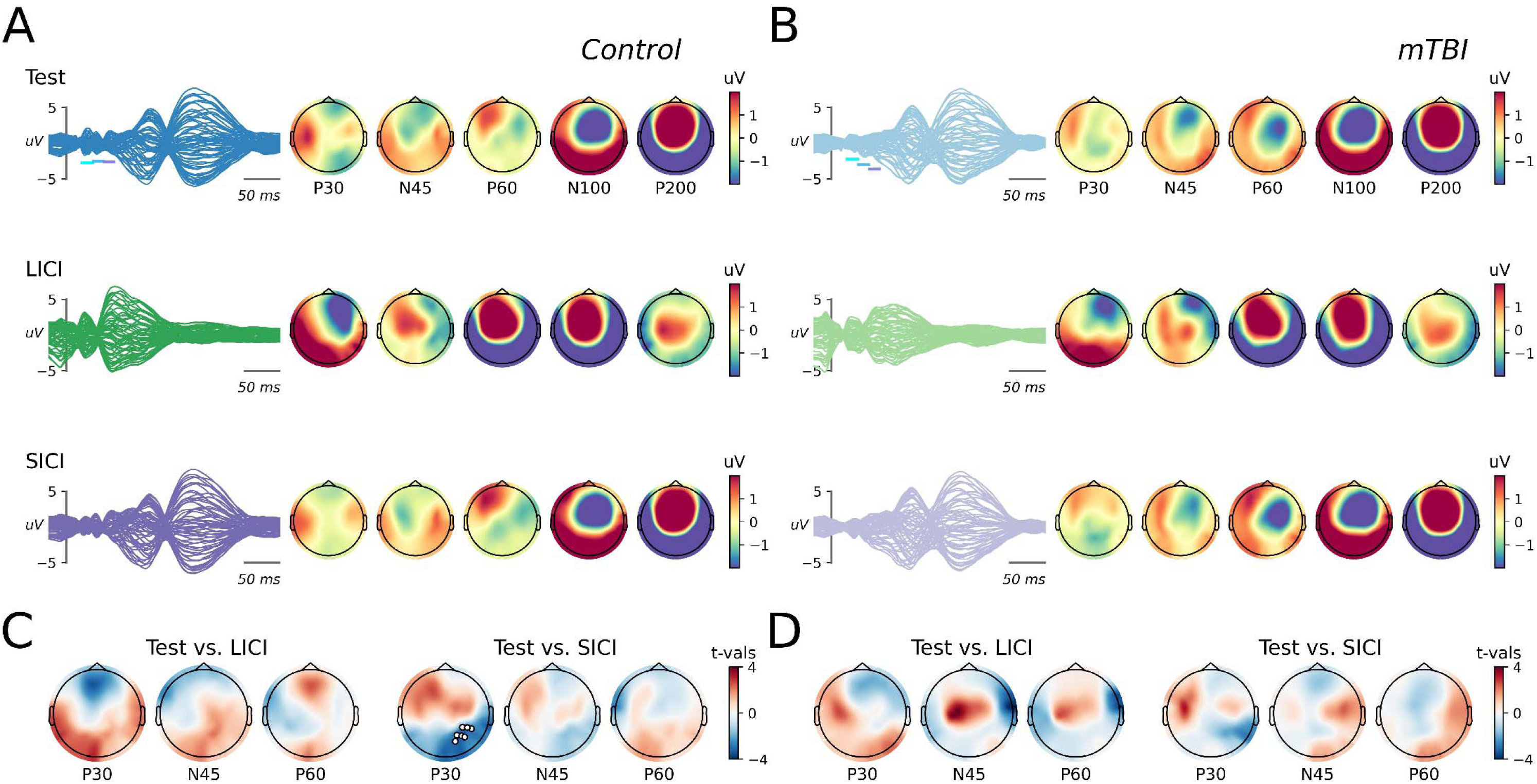
TEP response to ICI paradigms. **(A, B)** Butterfly plots and associated scalp topographies for canonical TEP components recorded following application of test-alone stimulation *(top)*, LICI *(middle)* or SICI *(bottom)* in healthy controls *(A)* and mTBI patients *(B)*. Coloured bars on butterfly plots indicate the time period over which each component was quantified. **(C, D)** For each component of interest, topoplots show results of within-group cluster-based comparisons between the response to single and paired-pulse stimulation in healthy controls *(C)* and mTBI patients *(D)*. White dots show electrodes within a s significant cluster.

**Figure 3.**
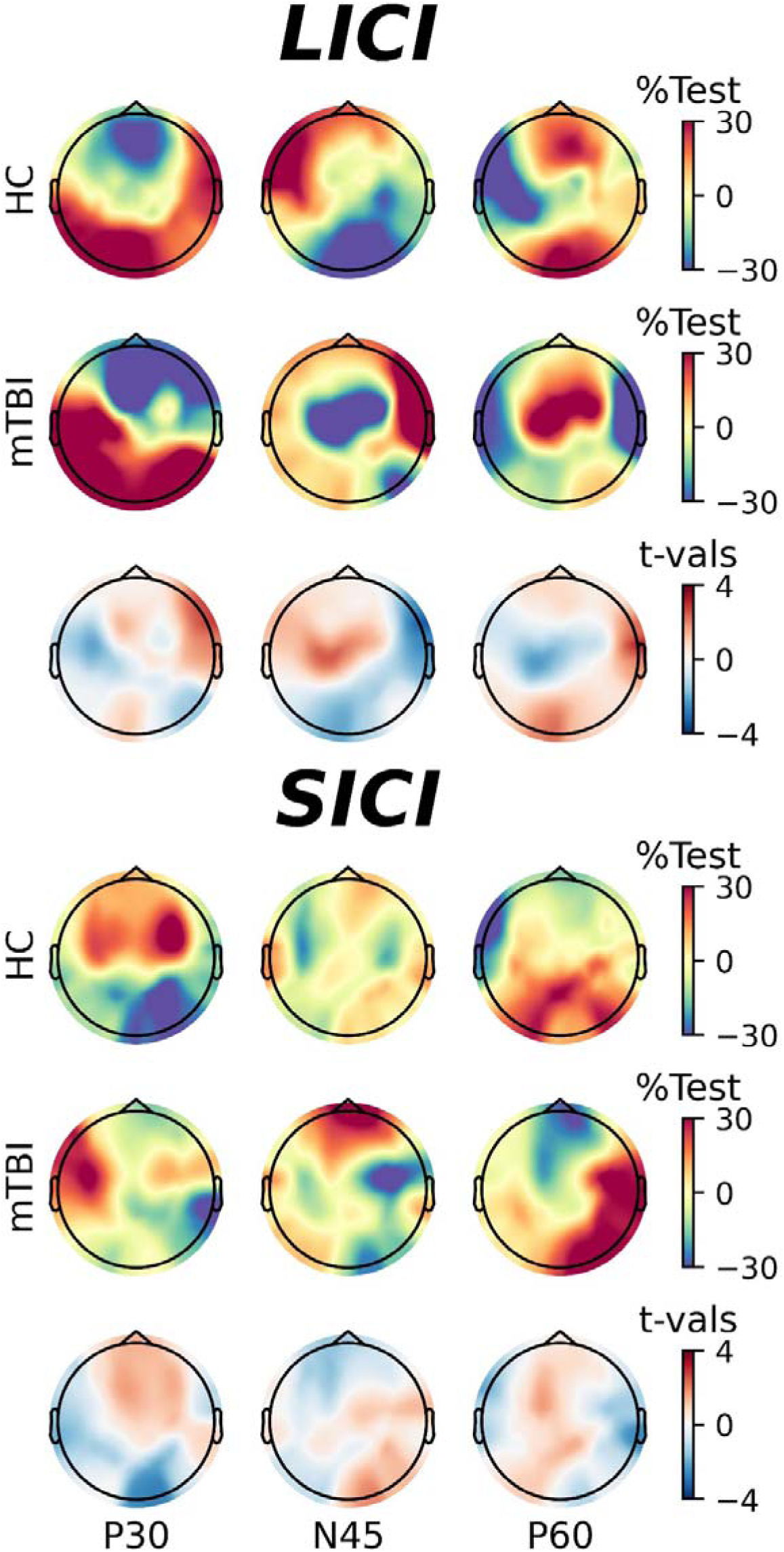
Effects of mTBI on TEP-measures of ICI. Normalised changes in TEP amplitude following application of LICI **(top panel)** and SICI **(bottom pane**l) in healthy controls *(top row)* and mTBI patients *(middle row)*, in addition to results of cluster-based comparisons between groups *(bottom row)*.

Figure 4 shows scalp topographies of TROs for the ICI session, separately for healthy controls (Fig 4A) and mTBI patients (Fig 4B), with results of within-subject comparisons shown in figure 4C/5D. For LICI, both groups showed reductions in oscillatory power in all bands (all *P*-values < 0.04). For SICI, the control group showed decreased power in theta (positive cluster: *P* = 0.004; positive cluster: *P* = 0.009), alpha (positive cluster: *P* = 0.0007; positive cluster: *P* = 0.02) and lower beta (positive cluster: *P* = 0.01) bands, in addition to increased power in the gamma band (negative cluster: *P* = 0.004)(Fig 4C). For mTBI patients, SICI was associated with reduced power in theta (positive cluster: *P* = 0.002; positive cluster: *P* = 0.03), alpha (positive cluster: *P* = 0.04) and upper beta (positive cluster: *P* = 0.01) bands (Fig 4D). Between-group comparisons of the normalised response to paired-pulse stimulation showed that effects of SICI on upper-beta activity were significantly greater in mTBI patients, consistent with an injury-related increase of inhibition of upper beta power by SICI (Fig 5).

**Figure 4.**
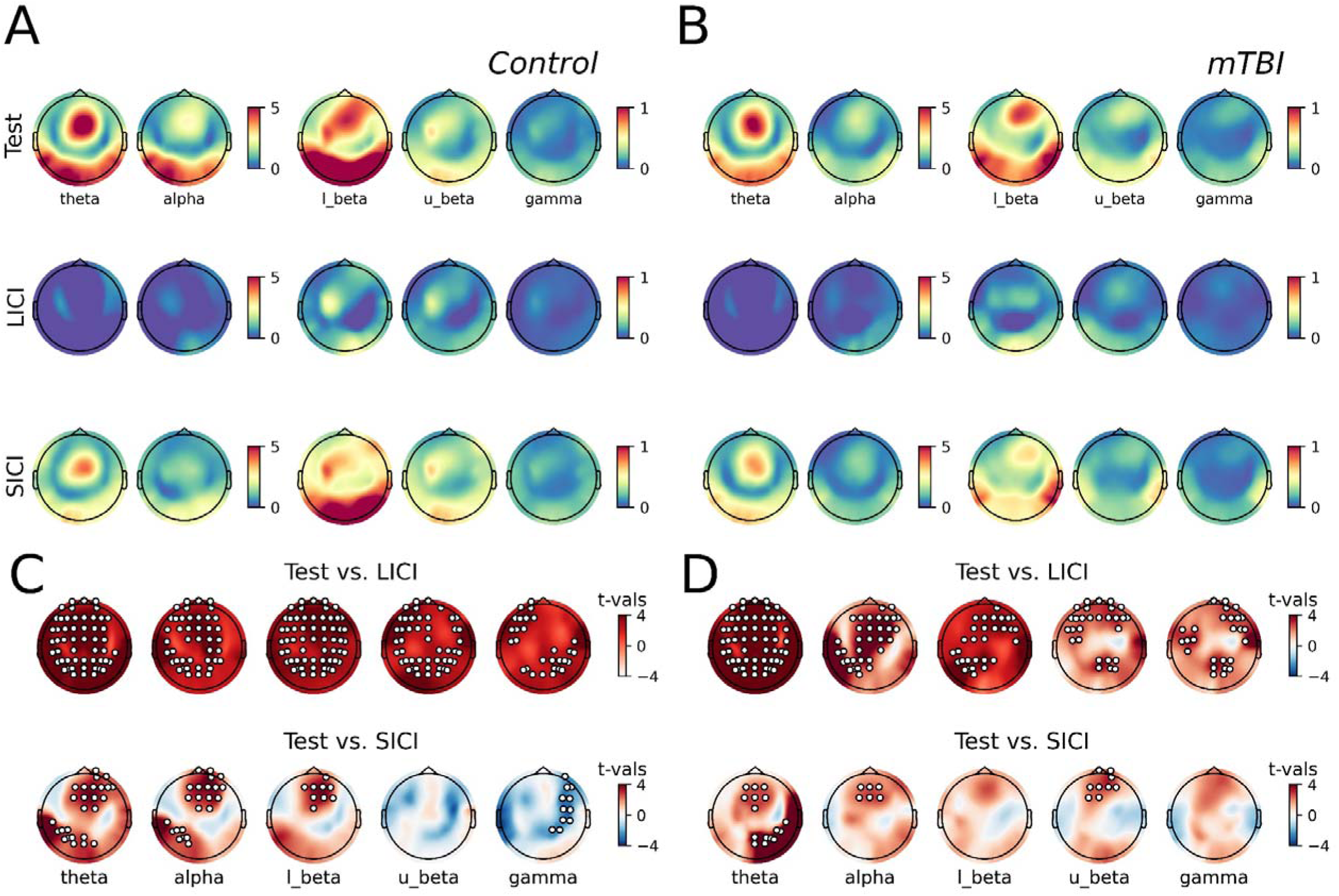
Response of TROs to ICI paradigms. **(A, B)** Plots of the time-frequency response beneath the coil (averaged over F3 and F5 electrodes) and associated scalp topographies for each band of interest following application of test-alone stimulation *(top)*, LICI *(middle)* or SICI *(bottom)* in healthy controls *(A)* and mTBI patients *(B)*. (**C, D**) For each band of interest, topoplots show results of within-group cluster-based comparisons between the response to single and paired-pulse stimulation in healthy controls *(C)* and mTBI patients *(D)*. White dots show electrodes within a s significant cluster.

**Figure 5.**
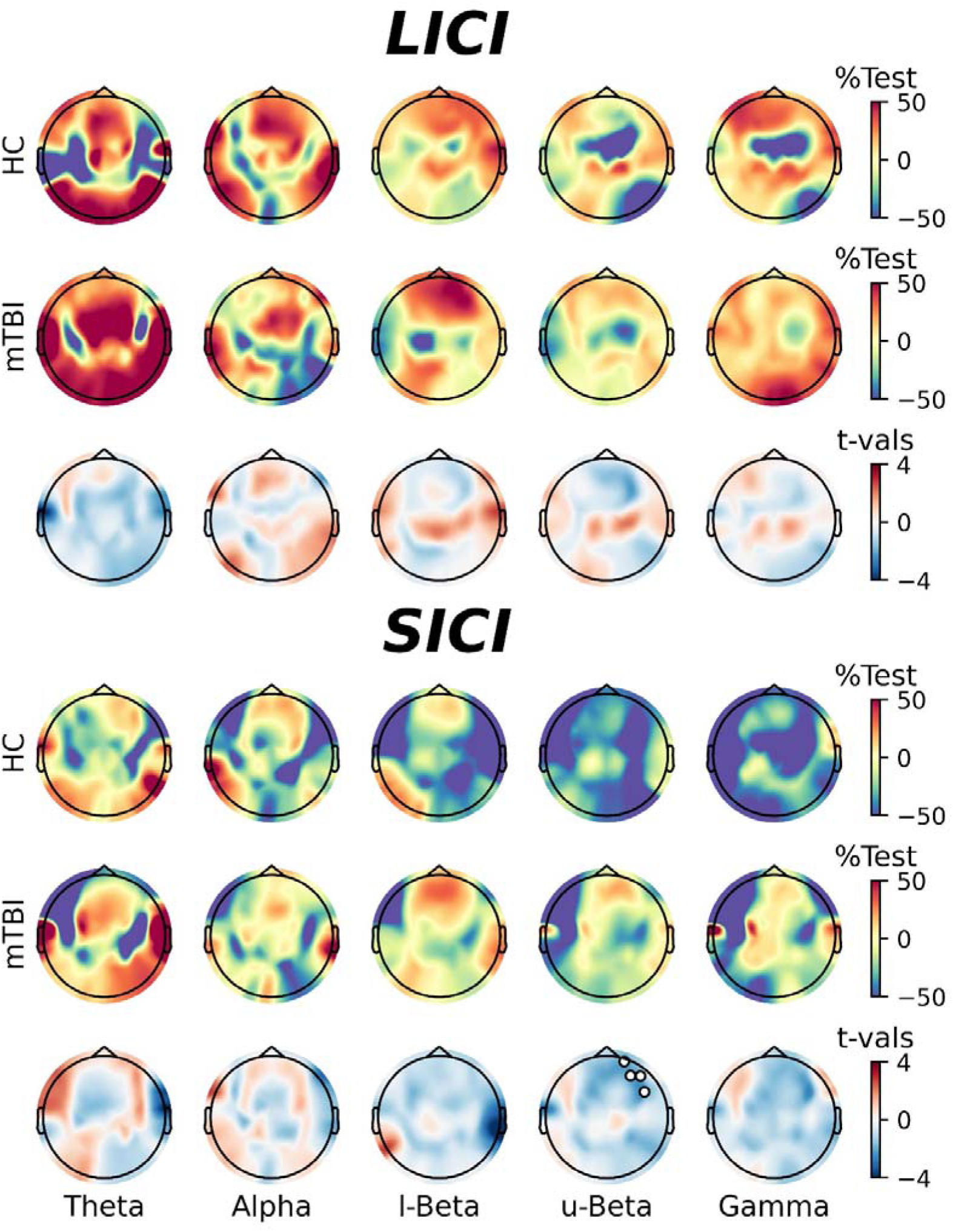
Effects of mTBI on the oscillatory indices of ICI. Normalised changes in TMS-related oscillations following application of LICI (**top panel**) and SICI (**bottom panel**) in healthy controls *(top row)* and mTBI patients *(middle row*), in addition to results of cluster-based comparisons between groups *(bottom row)*.

### cTBS

#### ROI analysis

Figure 6 shows results of ROI analyses from the cTBS session, including data from both time (Fig 6A/6B) and time-frequency (Fig 6C/6D) domains. For healthy controls at the Post 30 time point, there was a consistent (*pd* = 98.8%) and significant (0% in ROPE) increase in N45, as well as a consistent (*pd* = 98.8%) and significant (0% in ROPE) reduction in P60. Both groups also demonstrated consistent (*pd*: HC = 97.4%, mTBI = 95.4%) reductions in N45 at Post 0, but these failed to reach a practical level of significance (% in ROPE: HC = 1%, mTBI = 3%). Comparisons of normalised changes in N45 also revealed that the response of mTBI patients at Post 30 was consistently (*pd* = 95.3%) reduced relative to healthy controls, but this failed to reach a practical level of significance (0.9% in ROPE). All other comparisons in the time domain were inconsistent and failed to provide sufficient evidence to reject or accept the null hypothesis (all *pd* < 92.8%, % in ROPE: 6.7 – 35.0).

**Figure 6.**
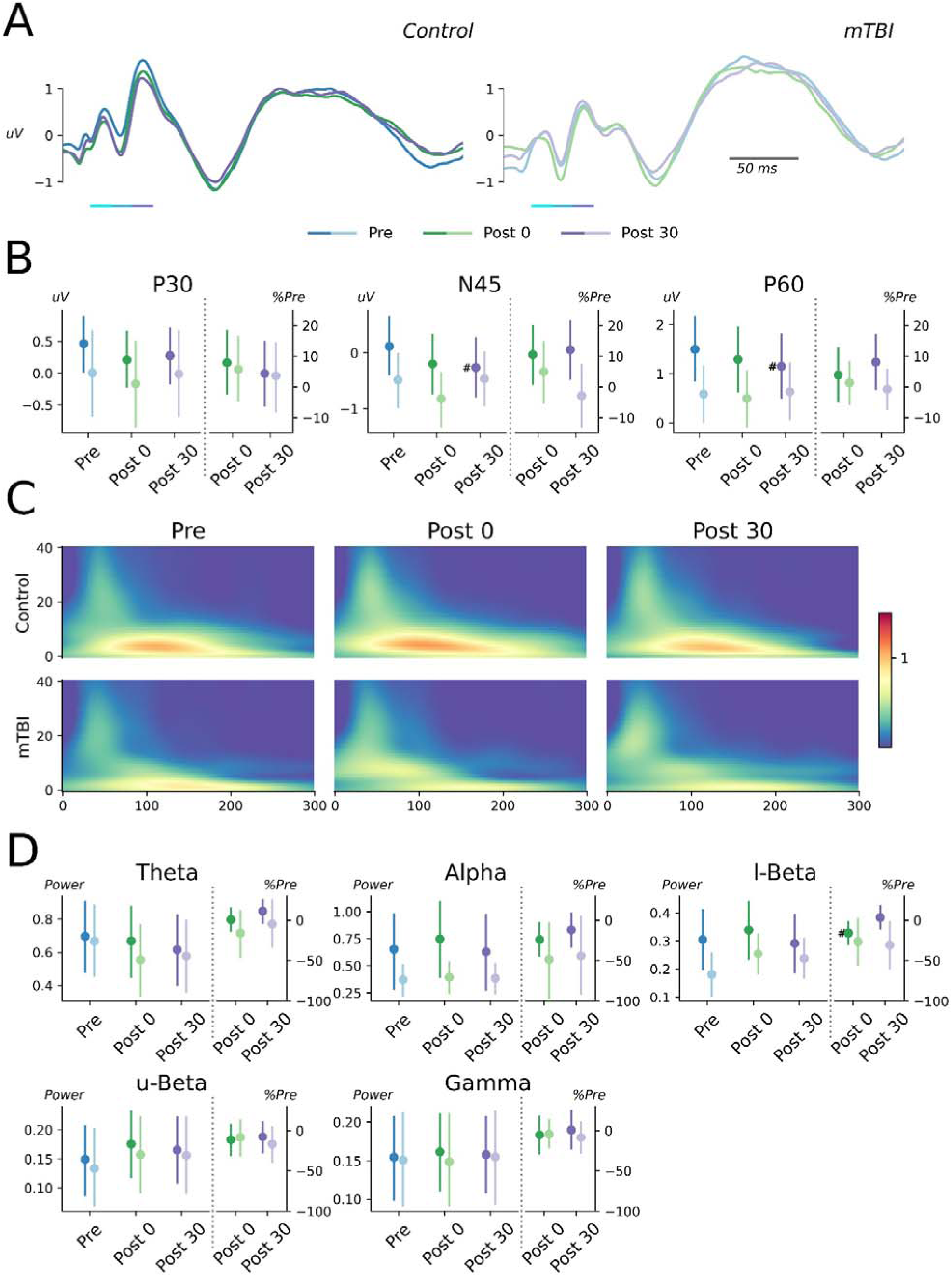
Results of ROI analysis from cTBS session. **(A)** TEPs from F3-F5 electrodes recorded at Pre *(blue line)*, Post 0 *(green line)* and Post 30 *(purple line)* time points in healthy controls *(left panel)* and mTBI patients *(right panel)*. Coloured bars below line plots highlight time periods used for quantification of TEP peaks. **(B)** Amplitude of P30 *(left panel)*, N45 *(middle panel)* and P60 *(right panel)* for each time points, compared between healthy controls *(dark colours)* and mTBI patients *(pale colours)*. Raw values are compared on the left y-axis, normalised values are compared on the right y-axis. **(C)** Time-frequency response at Pre *(left panels)*, Post 0 *(middle panels)* and Post 30 *(right panels)* time points for healthy controls *(top row)* and mTBI patients *(bottom row)*. **(D)** TRO power in theta, alpha, lower beta *(top row)*, upper beta and gamma *(bottom row)* bands for each time point, compared between healthy controls and mTBI patients. Raw values are compared on the left y-axis, normalised values are compared on the right y-axis. ^#^Significant and consistent difference relative to Pre/Post 0.

For TROs, healthy controls showed a greater normalised change in lower beta power at Post 0 compared to Post 30 which was consistent (*pd* = 98.3%) and significant (0% in ROPE). Furthermore, normalised changes in lower beta power at Post 30 were consistently (*pd* = 94.8%) greater in mTBI patients when compared to controls, but this failed to reach a practical level of significance (2% in ROPE). All other comparisons in the spectral domain were inconsistent and failed to provide sufficient evidence to reject or accept the null hypothesis (all *pd* < 88.4%, % in ROPE: 6.8 – 36.4).

#### Global analysis

Figure 7 shows grand average waveforms and scalp topographies for responses recorded at Pre, Post 0 and Post 30 time-points, in healthy controls (Fig 7A) and mTBI patients (Fig 7B). Within group comparisons failed to identify significant clusters for any comparison (Fig 7C, 7D). Between-group comparisons of normalised post-intervention responses also failed to identify any significant clusters (Fig 8).

**Figure 7.**
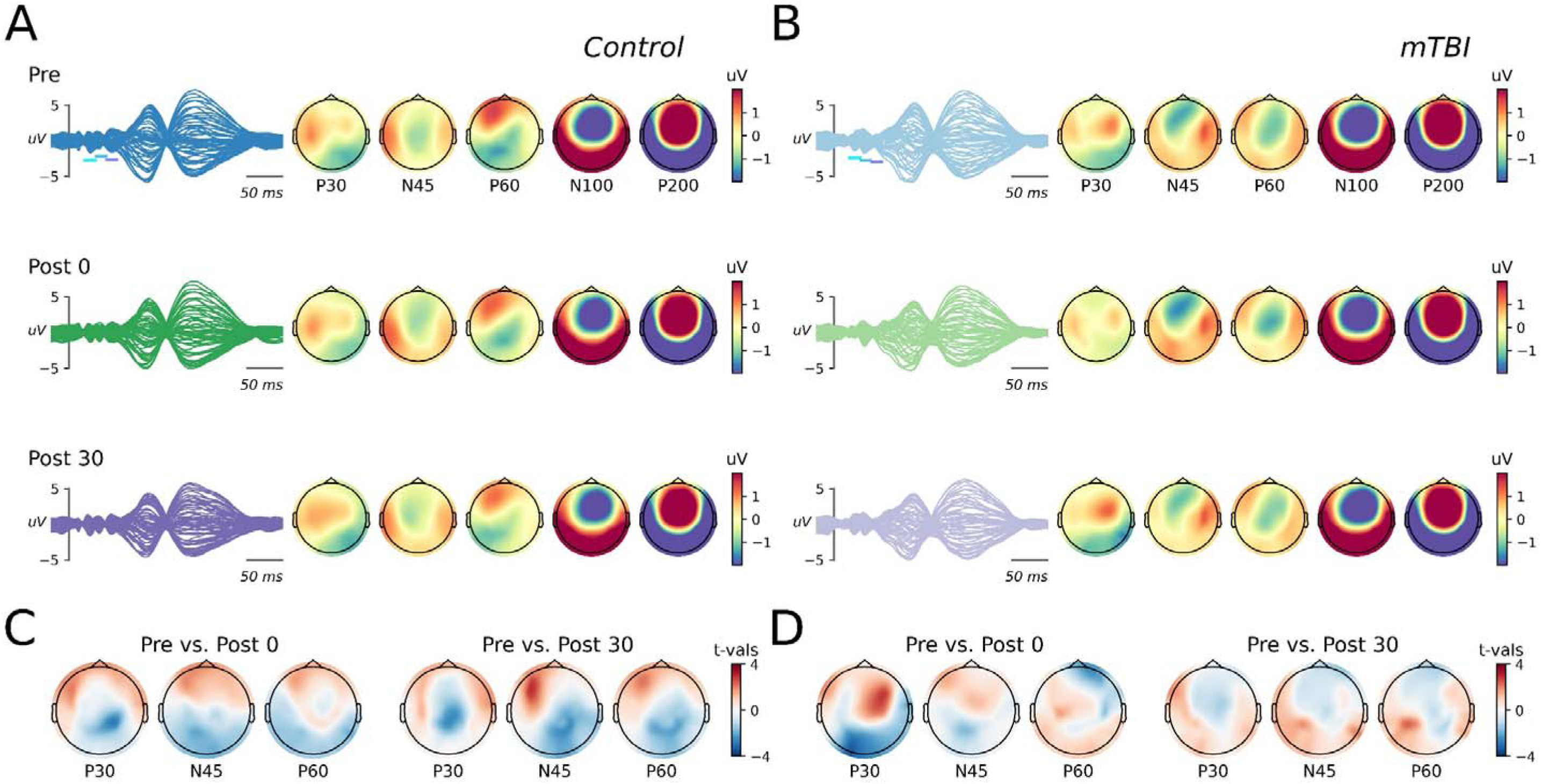
TEP response to cTBS. (**A, B**) Butterfly plots and associated scalp topographies for canonical TEP components recorded before (*Pre, top row*), immediately after (*P0, middle ro*w) and 30 mins after (*P30, bottom row*) application of cTBS in healthy controls *(A)* and mTBI patients *(B)*. Coloured bars on butterfly plots indicate the time period over which each component was quantified. (**C, D**) For each component of interest, topoplots show results of within-group cluster-based comparisons between pre- and post-intervention responses in healthy controls *(C)* and mTBI patients *(D)*.

**Figure 8.**
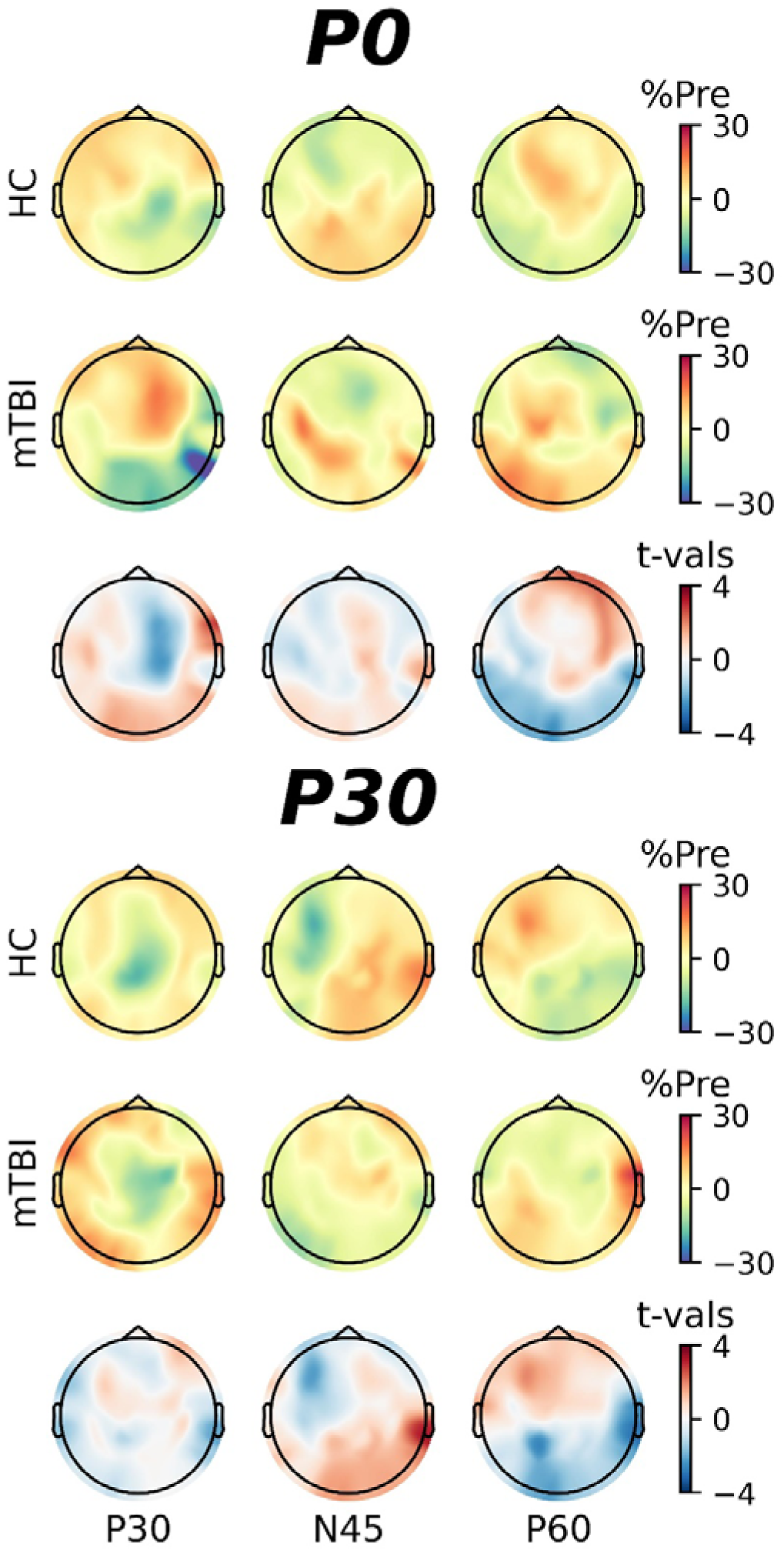
Effects of mTBI on response to cTBS. Normalised changes in TEP amplitude at P0 (**top panel**) and P30 (**bottom pane**l) time points in healthy controls *(top row)* and mTBI patients *(middle row)*, in addition to results of cluster-based comparisons between groups *(bottom row)*.

Figure 9 shows scalp topographies of TROs for the cTBS session, separately for healthy controls (Fig 9A) and mTBI patients (Fig 9B), with results of within-subject comparisons shown in figure 9C/9D. For the control group, power at the Post 30 timepoint was reduced in the theta (positive cluster: *P* = 0.01) and alpha (positive cluster: P = 0.04) bands (Fig 9C). For the mTBI group, power at the Post 0 timepoint was reduced in the upper beta (positive cluster: *P* = 0.04) and gamma (positive cluster: *P* = 0.02) bands, whereas power at the Post 30 timepoint was reduced in the theta (positive cluster: *P* = 0.03, positive cluster: *P* = 0.04), alpha (positive cluster: *P* = 0.03) and upper beta (positive cluster: *P* = 0.04) bands (Fig 9D). Between-group comparisons of normalised changes in oscillatory activity failed to identify any significant clusters (Fig 10).

**Figure 9.**
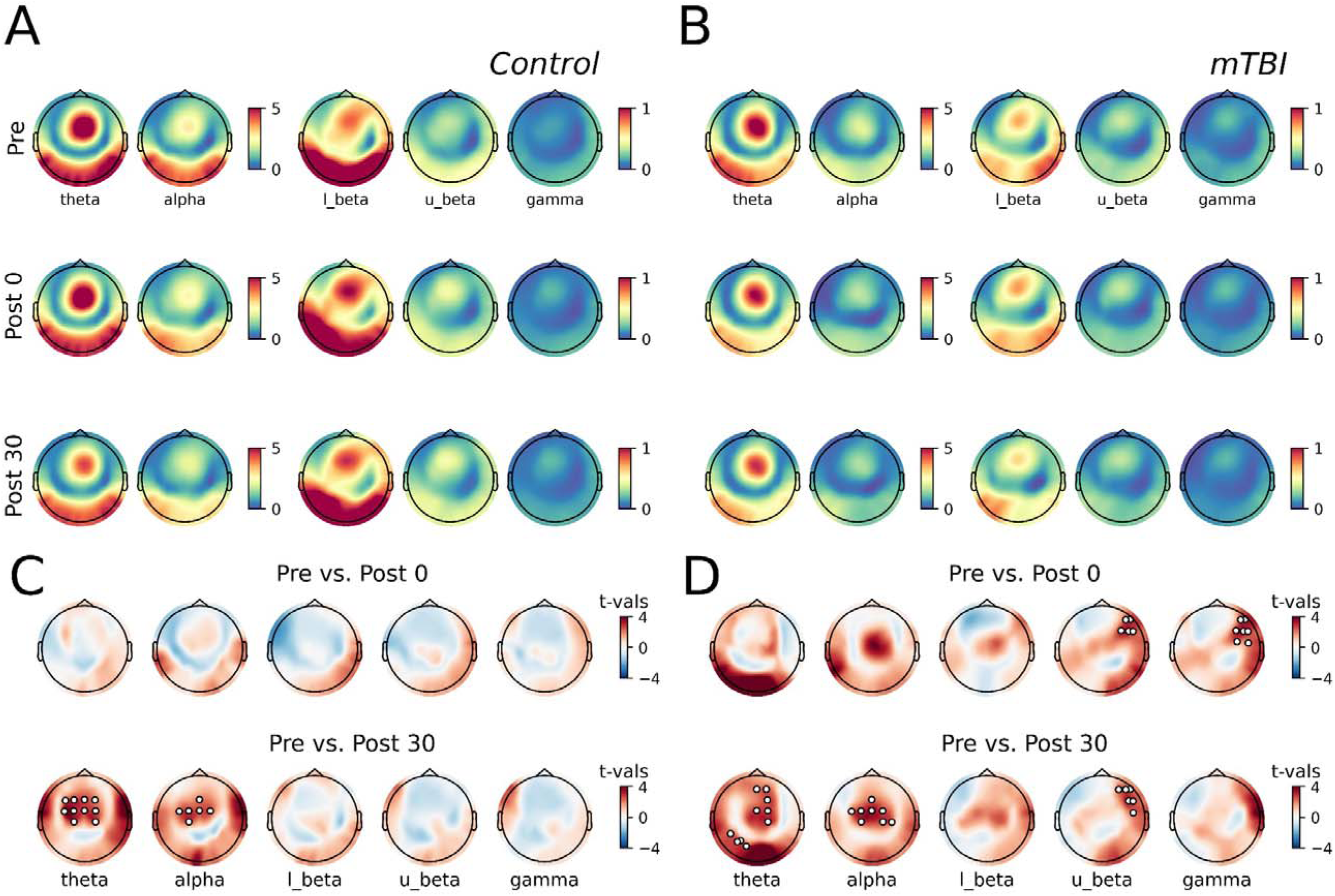
Response of TROs to application of cTBS. (**A, B**) Plots of the time-frequency response beneath the coil (averaged over F3 and F5 electrodes) and associated scalp topographies for each band of interest at Pre (*top*), P0 (*middle*) and P30 (*bottom*) timepoints in healthy controls *(A)* and mTBI patients *(B)*. (**C, D**) For each band of interest, topoplots show results of within-group cluster-based comparisons between pre- and post-intervention responses in healthy controls *(C)* and mTBI patients *(D)*. White dots show electrodes within a s significant cluster.

**Figure 10.**
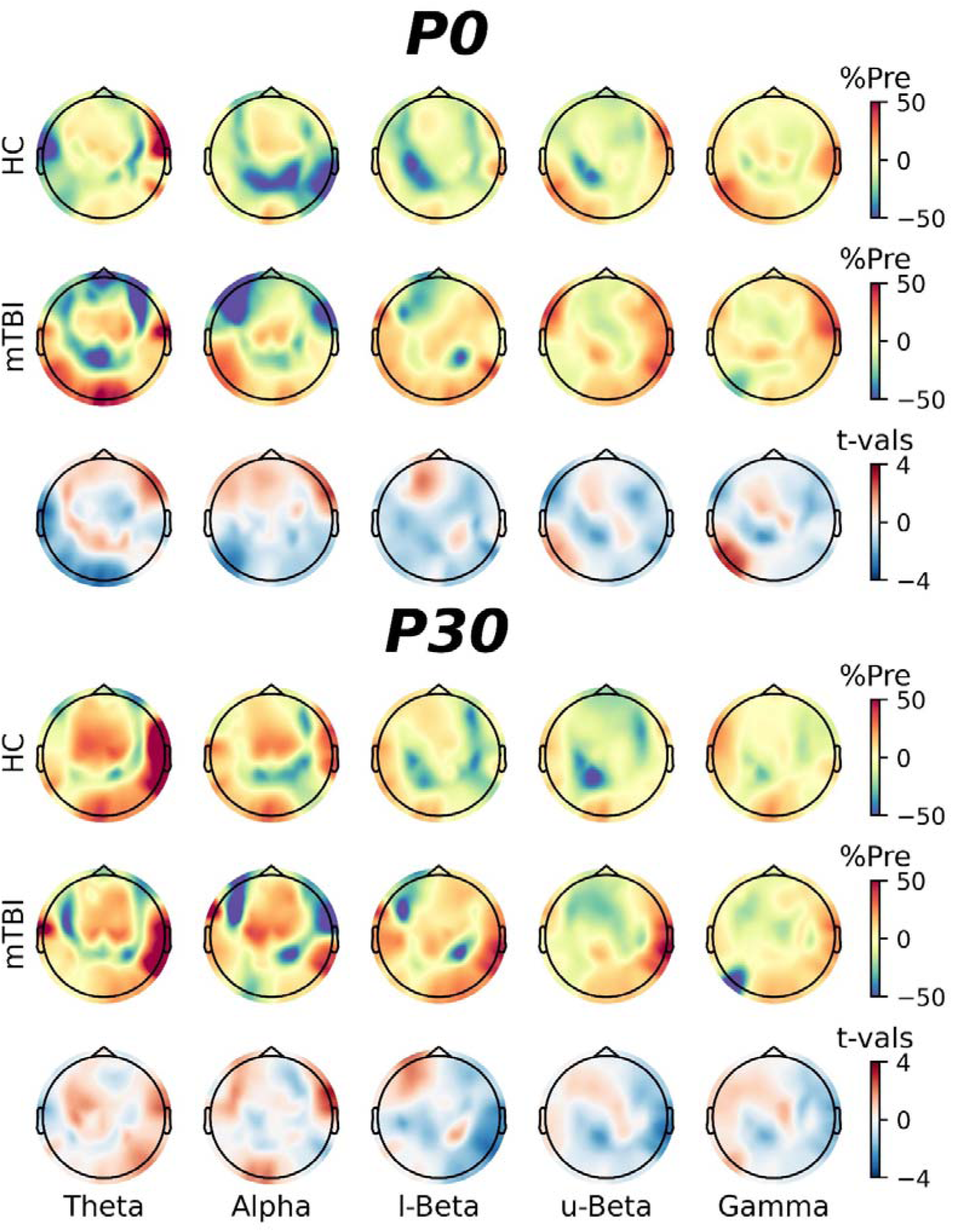
Effects of mTBI on the oscillatory response to cTBS. Normalised changes in TMS-related oscillations at P0 **(top panel)** and P30 **(bottom panel)** timepoints in healthy controls *(top row)* and mTBI patients *(middle row)*, in addition to results of cluster-based comparisons between groups *(bottom row)*.

### Executive function

Performance on each subtest of the *N*-back and TMT is compared between healthy controls and mTBI patients in Figure 11. For *N*-back, all within-group comparisons of *d*-prime and reaction time were inconsistent and failed to provide sufficient evidence to reject or accept the null hypothesis (all *pd* < 71.4%, % in ROPE: 11.4 – 12.6). For TMT-A, mTBI patients showed consistent (*pd* = 98.3%) and significant (0% in ROPE) reductions in accuracy compared with the control group. All other between-group comparisons were inconsistent and failed to provide sufficient evidence to accept or reject the null hypothesis (all *pd* < 87.1%, % in ROPE: 5.3 – 12.7).

**Figure 11.**
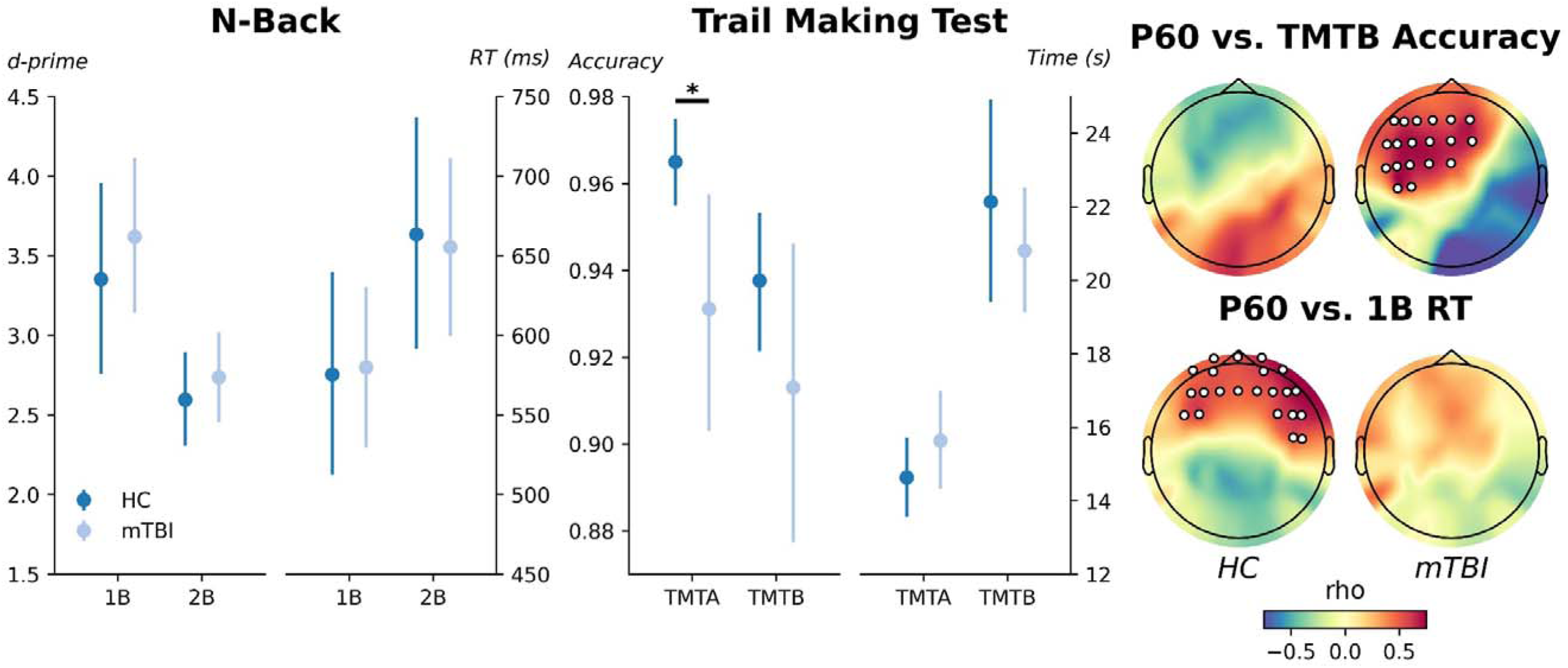
Effects of mTBI on executive function. (left panel) Performance on 1-back and 2-back variants of the N-back task, indexed via the d-prime metric and reaction time, compared between healthy controls *(dark blue circles)* and mTBI patients *(light blue circles)*. **(middle panel)**. Performance on the trail making test part A and B, indexed via accuracy and completion time, in healthy controls and mTBI patients. **(right panel)** Topographical distribution of significant cluster-based correlations between normalised changes in P60 amplitude 30 minutes after cTBS and TMT-B accuracy *(top row)* or 1-back reaction time *(bottom row)*. White circles show electrodes included in a significant cluster.

### Correlations

To investigate relationships between neurophysiological and functional effects of mTBI, TMS-EEG data were correlated with *N*-back, TMT scores and time-since-injury using cluster-corrected Spearman’s correlations. This identified that accuracy during the TMT-B subtest was related to normalised changes in P60 amplitude at Post 30 in the mTBI group (positive cluster: FDR-corrected *P* = 0.04) but not controls (Fig 11, right-upper panel). In contrast, reaction time in the 1-back test was related to normalised changes in P60 amplitude at Post 30 in controls (positive cluster: FDR-corrected *P* = 0.03) but not the mTBI group (Fig 11, right-lower panel). All other comparisons failed to identify any significant clusters.

## Discussion

The current study aimed to characterise effects of mTBI on intracortical inhibitory function and LTD-like neuroplasticity in DLPFC. This was achieved by using TMS-EEG applied over DLPFC to index SICI and LICI (assessing intracortical inhibition), as well as the response to cTBS (assessing LTD-like plasticity), in participants that had suffered an mTBI in the previous 2 years. These responses were then compared to a group of healthy control participants without a history of head injury. For analyses in the time domain (i.e., TEPs), application of both paired-pulse TMS and cTBS modulated early TEP amplitude, but this was only apparent in the healthy control group. In contrast, analyses of the spectral response to TMS (i.e., TROs) revealed modulatory effects of all paradigms in both groups. However, the mTBI group tended to show an increased inhibitory response in higher frequency bands.

Additionally, post-intervention responses to cTBS differentially predicted cognitive performance. Together these outcomes provide new evidence that mTBI is associated with neurophysiological changes within DLPFC.

### Response to SICI and LICI

Alterations to intracortical inhibitory function represent some of the most reported outcomes of studies applying TMS after mTBI. In particular, several conventional TMS studies have identified injury-related changes in the activity of intracortical circuits involving GABA_B_ receptors (De Beaumont *et al*. 2009, Tremblay *et al*. 2011, De Beaumont *et al*. 2012, Miller *et al*. 2014, Pearce *et al*. 2014, Di Virgilio *et al*. 2016, Edwards and Christie 2017), with some evidence also suggesting changes in GABA_A_-mediated inhibition (Pearce *et al*. 2014).

Importantly, this altered inhibitory function has been linked to mTBI severity (De Beaumont *et al*. 2012), demonstrating the potential clinical relevance of these neurophysiological alterations. While these observations are important, a significant limitation of the field has been that injury-related changes to specific TMS-based indices of inhibitory function (i.e., via L/SICI or the cortical silent period) have only been recorded from primary motor cortex. In contrast, non-motor areas of cortex are likely to be important in the cognitive effects of mTBI; quantifying how inhibitory function in these areas is influenced by injury therefore has the potential to better inform our understanding of how cognitive side-effects develop following mTBI.

Within the current study, the control group showed inhibition of P30 in response to SICI (for both ROI and global analyses), whereas LICI inhibited N45 (ROI analyses only), and these effects are consistent with findings from some previous studies (Rogasch *et al*. 2015, Desforges *et al*. 2022, Takano *et al*. 2025). While it is well established that conventional measures of SICI and LICI (i.e., those quantifying the response to stimulation with MEPs) reflect GABAergic processes, the mechanisms underpinning these measures when indexed with TMS-EEG remain unclear. However, it is important to note that a pharmacological study by Premoli *et al*. (2018) suggests that TMS-EEG indices of the response to paired-pulse stimulation are unlikely to be direct representations of their MEP-based counterpart.

Consequently, the current state of the literature cannot support a specific interpretation of the mechanisms contributing to TMS-EEG indices of SICI and LICI, and further investigation is required. Nonetheless, it seems reasonable to suggest that these paradigms modulate neural activity, and that the nature of this modulation differs between paradigms; this is demonstrated by both MEP- and TEP- based indices.

In contrast to the control group, the mTBI group failed to demonstrate any significant changes in TEP amplitude in response to either SICI or LICI, and this was consistent across local and global analyses. Consequently, our time-domain data suggest that mTBI is associated with a reduction in the modulatory effects of SICI and LICI within DLPFC. However, it’s important to note that normalised indices of inhibition failed to provide sufficient evidence for differences between groups. Within the ROI analysis, consistent differences were noted for LICI of the N45, but these failed to meet our predefined criteria for practical significance. Consequently, further work is needed to better elucidate the nature of these effects. One important factor to be considered in future studies is standardisation of test TEP amplitude. This is common practice in conventional paired-pulse TMS measures, as the amplitude of the test motor-evoked potential can influence the amount of inhibition recorded (Opie and Semmler 2014). While we found no statistical evidence for between group differences in the response to single-pulse stimulation (in either local or global analyses), the 89% HDIs for ROI data suggest that the test TEP for all peaks of interest varied over a relatively broad range (P30: −0.3 to 1.6 µV; N45: −1.1 to 1.2 µV; P60: 0.05 to 2.4 µV), which would have increased variability of the normalised data. Standardisation of response amplitude was omitted from the current study as data collection began prior to the required tools being readily available. Toolboxes for adjusting stimulation parameters online to optimise TEPs using real-time feedback have recently been released and could be used in future studies (i.e., Casarotto *et al*. 2022).

To further explore the neurophysiological effects of mTBI, the spectral response to stimulation was also examined. In healthy controls, application of LICI was associated with inhibition of all oscillatory bands, and this was apparent in both ROI and global analyses. This pattern of response is consistent with outcomes reported by several previous studies (Farzan *et al*. 2009, Garcia Dominguez *et al*. 2014, Rogasch *et al*. 2015, Desforges *et al*. 2022), possibly reflecting generalised inhibition of DLPFC output by LICIs activation of GABA_B_-mediated processes, although pharmacological validation of the underlying mechanisms is currently missing (Rogasch *et al*. 2015). For the mTBI group, effects were consistent with the control group at the global level, but failed to include the alpha and lower beta bands in ROI analyses. However, there were no between-group differences in normalised LICI, locally or globally, for any band. Consequently, differences in ROI analyses may be more reflective of slight variability in the topographic distribution of the response to stimulation in these sensor-level recordings, as opposed to specific effects of mTBI. Future work implementing comparisons of spatially filtered data will be important for clarification here. Nonetheless, when taken together, the data suggests that the inhibitory response to LICI within the spectral domain is maintained in DLPFC after mTBI. While this may appear contradictory to results in the time domain, it is important to note that our spectral analysis quantified total power. This metric includes inputs from both event-locked (i.e., those predominantly influencing time domain responses) and non-event-locked processes (Roach and Mathalon 2008), which may explain the disparity between temporal and spectral results.

For SICI, ROI analyses in both groups failed to identify significant changes in any band. For the control group, this contrasts findings from Desforges *et al*. (2022), who reported inhibition across all frequency bands, and Cash *et al*. (2016), who reported inhibition specific to the alpha band. However, it seems likely that the disparity here stems from the previous studies including larger ROIs (Cash: 10 electrodes; Desforges: 5 electrodes) that can be expected to incorporate changes that extend more broadly across the scalp. In support of this, global analyses in the current study showed that the control group demonstrated inhibition of low frequency activity (i.e., theta, alpha and lower beta bands), but facilitation of high frequency activity (i.e., gamma band). As all previous investigations of the TMS-EEG response to SICI have focussed on ROI analyses, it is difficult to contextualise these results. However, recent work has suggested there may be some disconnect between the information reflected by low (i.e., < 12 Hz) and high (i.e., > 13Hz) frequency oscillatory activity generated by TMS (Biabani *et al*. 2021). Specifically, lower frequency oscillations possibly reflect neural activity relating to somatosensory input, whereas higher frequency oscillations are more reflective of a direct neural response to TMS (Biabani *et al*. 2021). In contrast to the previous work using ROI analyses in healthy participants, our global analyses would therefore suggest that application of SICI results in a facilitation of neural output. While we note the absence of evidence supporting a role of GABA in the effects of SICI within the DLPFC, this outcome is nonetheless consistent with previous work in both animals (Hofmann *et al*. 2019) and humans (Lozano-Soldevilla *et al*. 2014) that reported an increase in gamma power following pharmacological potentiation of GABA_A_ activity. Despite this, SICI tended to produce inhibition of higher frequency activity in the mTBI group, and this was significant for the upper beta band in both within- and between-group comparisons. Consequently, our results provide novel evidence that mTBI alters the spectral effects of SICI when applied to DLPFC. It will be interesting to validate this finding in an independent sample of mTBI patients, and to explore potential functional implications.

### Response to cTBS

For the control group, global analyses of the response to cTBS failed to identify any significant change in TEP amplitude. However, ROI analyses instead found significant potentiation of N45 and inhibition of P60. This outcome contrasts findings from two previous studies that failed to identify changes in the early TEP components following cTBS over DLPFC, both in ROI (Chung *et al*. 2017, Nikolin *et al*. 2025) and global analyses (Chung *et al*. 2017). While it remains unclear why the current study was able to identify significant changes in the early TEPs that were absent in previous work, methodological inconsistencies may have contributed, including differences in TBS intensity (current study: 70% RMT, Chung et al: 80% RMT, Nikolin et al: 75%RMT) and location of stimulation (current study: F3-F5 boundary, Chung et al: F3, Nikolin et al: F3). While N45 and P60 have been associated with GABAergic (Premoli *et al*. 2014, Darmani *et al*. 2016) and glutamatergic (Belardinelli *et al*. 2021) processes, respectively, it is important to note that this relationship has been established in M1. In contrast, some evidence suggests responses in DLPFC may involve different mechanisms (Rogasch *et al*. 2020), although it remains unclear what these are.

Nonetheless, there is consensus that the early TEP peaks are likely to reflect net activity of neural circuits activated by TMS (both excitatory and inhibitory), and the effects in the control group are therefore consistent with inhibitory effects of cTBS on neural activity. In contrast, the mTBI group failed to show any change in early TEP amplitude following cTBS. Furthermore, normalised changes in N45 were consistently reduced in patients, only just failing to reach a practical level of significance (<1% in ROPE). Consequently, our time-domain data suggest a reduced response of DLPFC to cTBS following mTBI, particularly for the N45 potential.

For data in the spectral domain, ROI analyses failed to show any significant effects in either group. However, global analyses in controls revealed significant inhibition of activity in theta and alpha bands – a response that is partially consistent with Chung *et al*. (2017), who reported a reduction in ROI theta power after cTBS. While a similar response was also apparent in the mTBI group, they additionally demonstrated reductions of upper beta and gamma bands. Although the cause of this response is currently unclear, it is interesting to note the similarities with the spectral response to SICI (i.e., increased inhibitory response specific to high-frequency bands). Given the importance of GABA_A_-mediated processes in generating both SICI (Ziemann *et al*. 1996) and high-frequency oscillatory activity (Traub *et al*. 1996), it may therefore be possible that these similar spectral responses to separate interventions are underpinned by a common neurophysiological effect of mTBI (i.e., GABAergic dysregulation). This supports conclusions from our previous TMS-EEG work in primary motor cortex of mTBI patients, which also identified greater changes in GABA_A_ function (Opie *et al*. 2019).

### Interaction between function and neurophysiology

While N-back performance was not different between groups, patients showed slightly lower accuracy in TMT-A, demonstrating a subtle injury-related deficit in visual attention and executive function (Reitan 1958). However, this was unrelated to any neurophysiological measures. In contrast, normalised changes in P60 amplitude recorded 30 mins after cTBS were found to have differential relationships with performance in patients and controls.

Specifically, a larger reduction in responses tended to correlate with increased TMT-B accuracy in mTBI patients but not controls. In contrast, more facilitated responses tended to correlate with increased reaction times on the 1-back for controls but not patients. Specific interpretation of these results is challenging, particularly given they appear unrelated to the studies main outcomes. Nonetheless, they appear to suggest that, irrespective of group, reduction of P60 following cTBS is beneficial for performance. This is consistent with previous work in older individuals, where increases in the P60 following iTBS to left DLPFC were correlated with improved performance on a paired-associative learning task (Goldsworthy et al, 2020), suggesting that preserved plasticity in the prefrontal cortex is important for cognitive performance. While the mechanisms that underpin P60 remain unclear, previous work has established connections with sensory reafference (Petrichella *et al*. 2017); while this was within the context of the motor system, it may be possible that it serves a similar role in afferent communication in non-motor areas. For example, increased inhibition of competing or distracting information could be beneficial in the context of the tasks completed here. Despite this, the differential response between groups could suggest an injury-related alteration to the nature of this relationship. However, this will require substantiation in future work.

In conclusion, the current study used TMS-EEG – in conjunction with paired-pulse TMS, cTBS, and tests of executive function – to investigate the physiological and functional effects of mTBI on DLPFC. Our results in the time domain were generally suggestive of reduced response in patients. In contrast, data in the spectral domain suggested an increased response in patients. These disparities highlight the potential importance of considering different facets of the complex activity indexed by EEG (i.e., induced and evoked activity) when characterising the neurophysiological effects of mTBI. Finally, although unrelated to specific neurophysiological effects of mTBI, interactions between P60 modulation with cTBS and executive function were identified. These highlight new avenues of investigation for future studies in the field.

## Supporting information

supp_results

## Data availability

The data that support the findings of this study are available on request from the corresponding author. The data are not publicly available due to privacy or ethical restrictions.

## Funding

This study was supported by funding from the Neurosurgical Research Foundation of South Australia. GMO was supported by funding from the National Healthy and Medical Research Council (APP1139723) and the Australian Research Council (DE230100022)

## Conflict of Interest

The authors have no conflicts to declare.

## Notes

### Competing Interest Statement

The authors have declared no competing interest.

