## Supplementary material for "Mild traumatic brain injury alters function in the dorsolateral prefrontal cortex: a TMS-EEG study": supp_results

### Supplementary Results

<u>Analysis of non-paired pulse corrected data</u>

*ROI analysis:*

Figure S1 shows results of ROI analyses from the ICI session, including data form both time (Fig S1A/S1B) and time-frequency (Fig S1C/S1D) domains. For P30, LICI was associated with a consistent reduction in mTBI patients (*pd* = 95.5%), but this failed to reach a practical level of significance (1.3% in ROPE). Between-group comparisons of normalised changes in P30 also showed consistent differences between groups (*pd* = 95.1%) that failed to reach a practical level of significance (1.2% in ROPE). For N45 in healthy controls, LICI was associated with a consistent (*pd* = 97.4%) and significant (0% in ROPE) reduction in amplitude, relative to test alone. For P60 in mTBI patients, LICI was associated with an increase in amplitude which was consistent (*pd* = 96.7%) and significant (0% in ROPE). Healthy controls also showed consistent increases in P60 with LICI (*pd* = 94.4%), but this failed to reach a practical level of significance (4.4% in ROPE). All other comparisons in the time domain were inconsistent and failed to provide sufficient evidence to reject or accept the null hypothesis (all *pd* < 90.2%, % in ROPE: 3.3 – 18.1).

For TMS-related oscillatory activity, application of LICI was associated with reduced power relative to test alone in alpha, lower and upper beta, and gamma bands for healthy controls and this was consistent (all *pd* > 97.6%) and significant (0% in ROPE for all comparisons). LICI in mTBI patients was also associated with consistent (*pd* > 99.7%) and significant (0% in ROPE) reduction in upper beta power. A consistent reduction in gamma power was also seen for LICI in mTBI patients (*pd* = 95.3%), but this failed to reach a practical level of significance (3.1% in ROPE). All other comparisons in the spectral domain were inconsistent and failed to provide sufficient evidence to reject or accept the null hypothesis (all *pd* < 89.5%, % in ROPE: 8.3 – 33.6).

*Global analysis:*

Figure S2 shows grand average waveforms and associated scalp topographies for each stimulus condition from the ICI session, separately for mTBI patients (Fig S1A) and healthy controls (Fig S1B). Relative to the Test response, application of LICI in the control group increased P30 (negative cluster: $P = 0.006$, positive cluster: $P = 0.009$), N45 (negative cluster: $P = 0.01$) and P60 (negative cluster: $P < 0.0001$, positive cluster: $P < 0.0001$)(Fig S2C). For the mTBI group, LICI produced potentiation of both P30 (negative cluster: $P = 0.003$, positive cluster: $P = 0.02$) and P60 (negative cluster: $P = 0.0003$, positive cluster: $P = 0.0004$)(Fig S2D). Between-group comparisons of normalised LICI and SICI values in the time domain failed to identify any significant clusters (Fig S3).

Figure S4 shows scalp topographies of TMS-related oscillatory activity for the ICI session, separately for healthy controls (Fig S4A) and mTBI patients (Fig S4B), with results of within-subject comparisons shown in figure S4C/S4D. Relative to Test, application of LICI in the both groups reduced power in all bands (all *P*-values < 0.04). In contrast, application of SICI reduced upper beta (negative cluster: $P = 0.03$) and gamma (negative cluster: $P = 0.0009$) power in healthy controls (Fig S4C), but reduced theta power in mTBI patients (negative cluster: $P = 0.01$) (Fig S4D). Between-group comparisons of normalised oscillatory activity failed to identify any significant clusters (Fig S5).

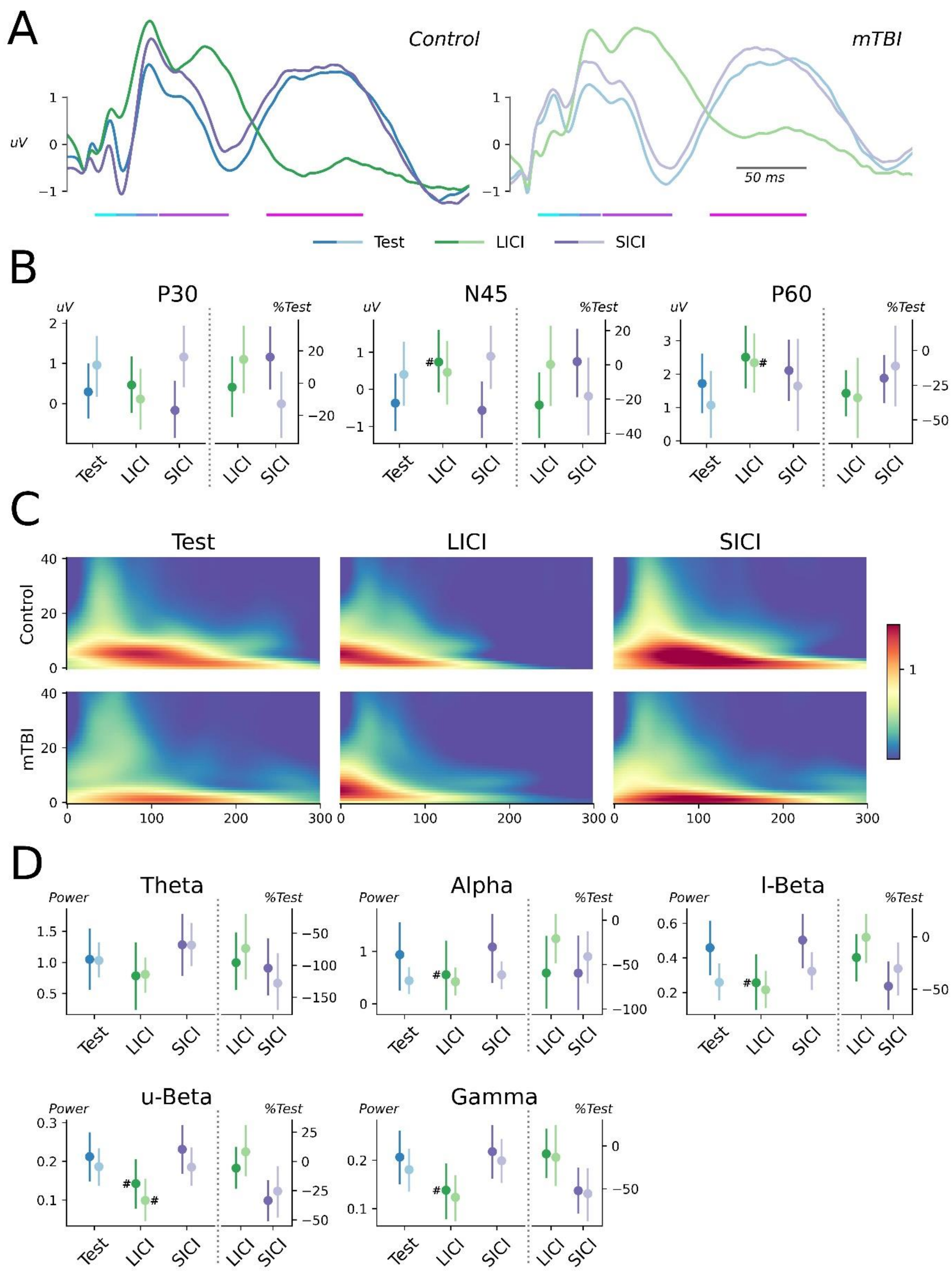


**Figure S1. Results of ROI analysis from ICI session without paired-pulse correction.** Data in all panels have not undergone the subtraction procedure to remove the TEP generated by the conditioning stimulus. **(A)** TEPs from F3-F5 electrodes recorded in response to test alone *(blue line)*, LICI *(green line)* and SICI *(purple line)* in healthy controls *(left panel)* and mTBI patients *(right panel)*. Coloured bars below line plots highlight time periods used for quantification of TEP peaks. **(B)** Amplitude of P30 *(left panel)*, N45 *(middle panel)* and P60 *(right panel)* for each stimulus condition, compared between healthy controls *(dark colours)* and mTBI patients *(pale colours)*. Raw values are compared on the left y-axis, normalised values are compared on the right y-axis **(C)** Time-frequency responses generated by application of test alone *(left panels)*, LICI *(middle panels)* and SICI *(right panels)* for healthy controls *(top row)* and mTBI patients *(bottom row)*. **(D)** TMS-related oscillatory power in theta, alpha, lower beta *(top row)*, upper beta and gamma *(bottom row)* bands for each stimulus condition, compared between healthy controls and mTBI patients. Raw values are compared on the left y-axis, normalised values are compared on the right y-axis. #Significant and consistent difference relative to test alone.

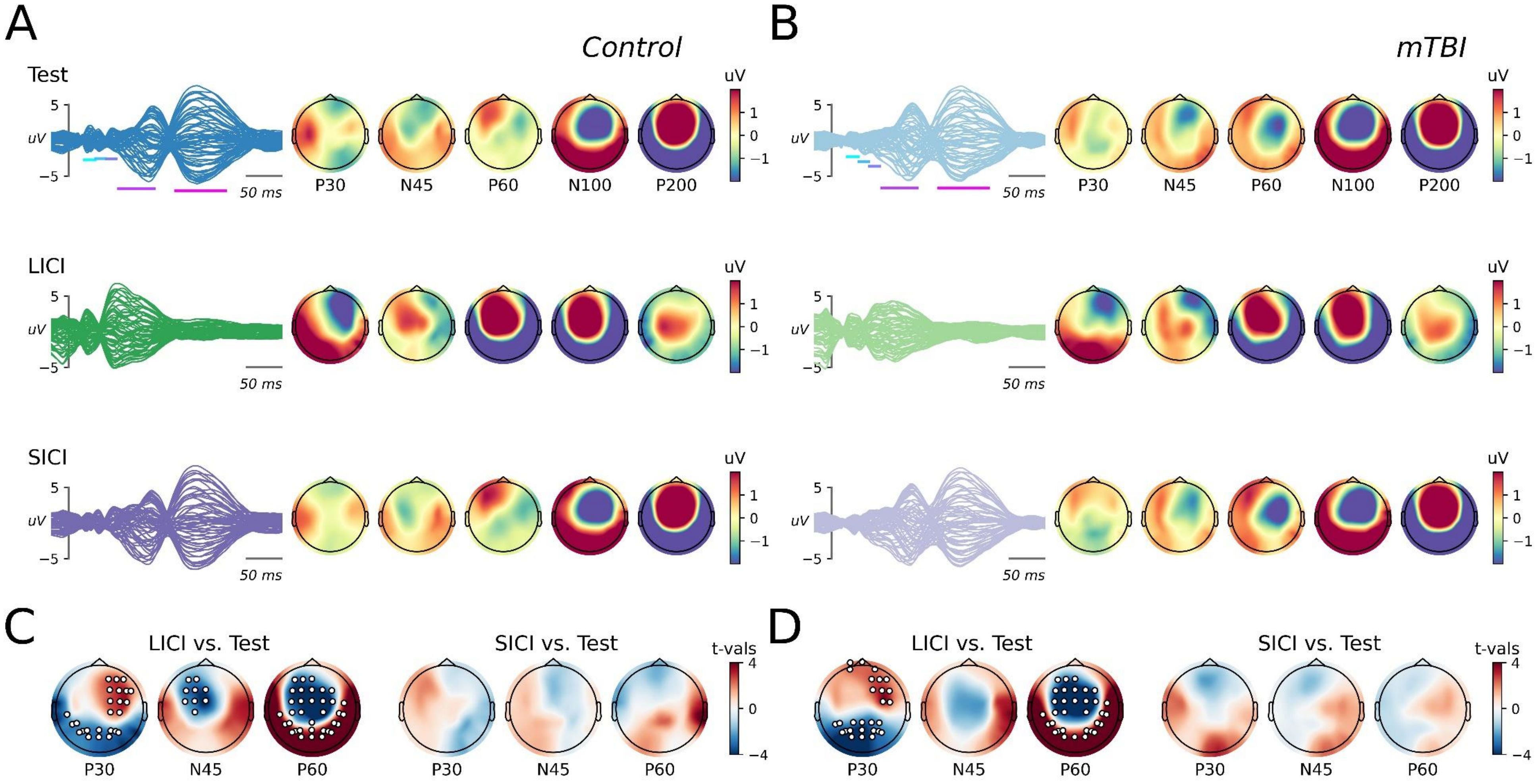


**Figure S2. TEP data from ICI paradigms without paired-pulse correction. (A, B)** Butterfly plots and associated scalp topographies for canonical TEP components recorded following application of test-alone stimulation *(top)*, LICI *(middle)* or SICI *(bottom)* in healthy controls *(A)* and mTBI patients *(B)*. Data have not undergone the subtraction procedure to remove the TEP generated by the conditioning stimulus. Coloured bars on butterfly plots indicate the period over which each component was quantified. **(C, D)** For each component of interest, topoplots show results of within-group cluster-based comparisons between the response to single and paired-pulse stimulation in healthy controls *(C)* and mTBI patients *(D)*. White dots show electrodes within a s significant cluster.

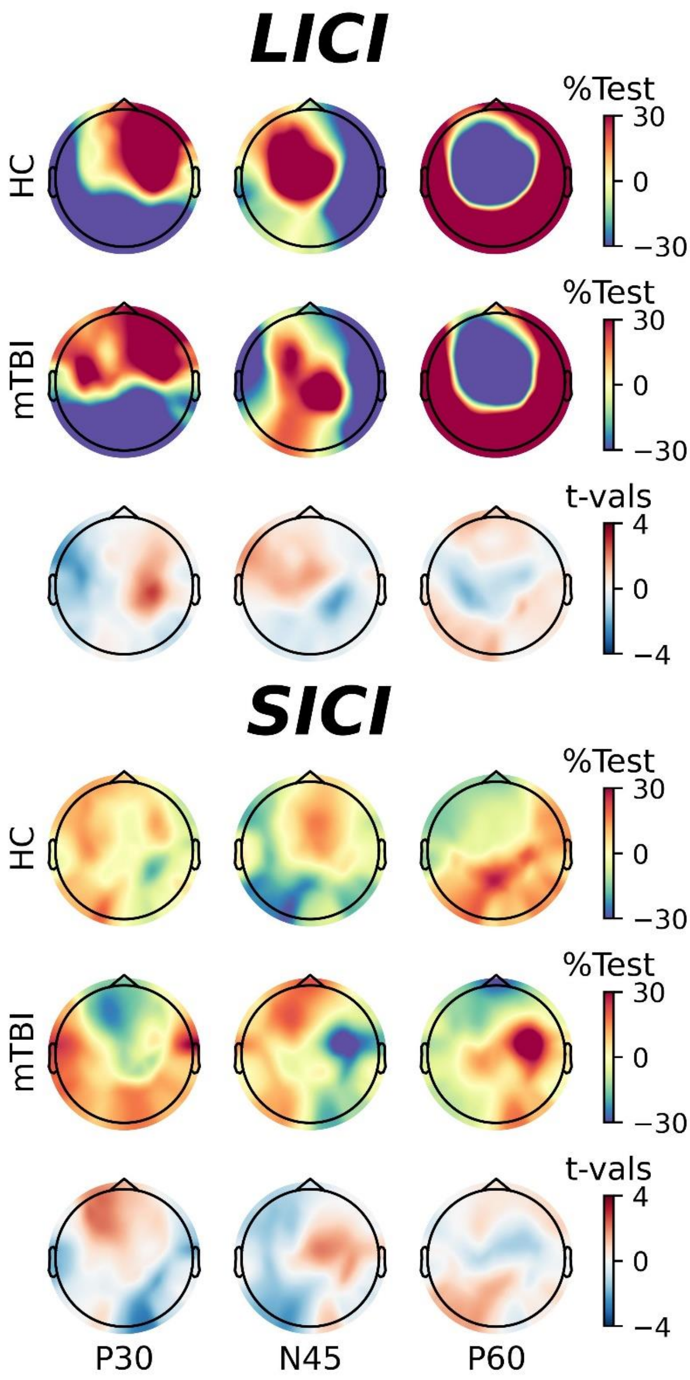


**Figure S3. Effects of mTBI on TEP-measures of ICI without paired-pulse correction.** Normalised changes in TEP amplitude following application of LICI **(top panel)** and SICI **(bottom panel)** in healthy controls *(top row)* and mTBI patients *(middle row)*, in addition to results of cluster-based comparisons between groups *(bottom row)*. Data have not undergone the subtraction procedure to remove the TEP generated by the conditioning stimulus.

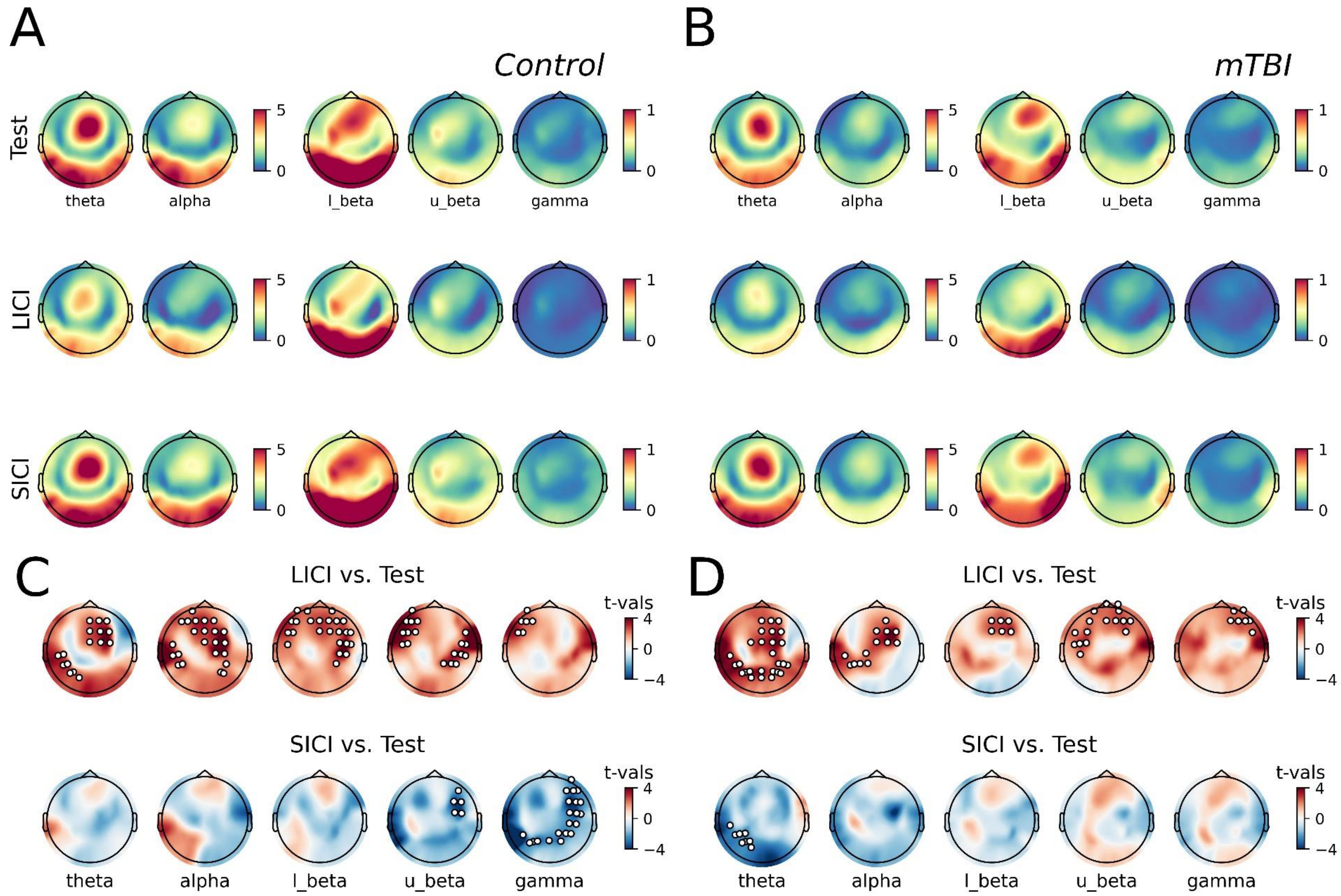


**Figure S4. Response of TMS-related oscillatory activity to ICI paradigms without paired-pulse correction. (A, B)** Plots of the time-frequency response beneath the coil (averaged over F3 and F5 electrodes) and associated scalp topographies for each band of interest following application of test-alone stimulation *(top)*, LICI *(middle)* or SICI *(bottom)* in healthy controls *(A)* and mTBI patients *(B)*. Data have not undergone the subtraction procedure to remove the TEP generated by the conditioning stimulus. **(C, D)** For each band of interest, topoplots show results of within-group cluster-based comparisons between the response to single and paired-pulse stimulation in healthy controls *(C)* and mTBI patients *(D)*. White dots show electrodes within a s significant cluster.

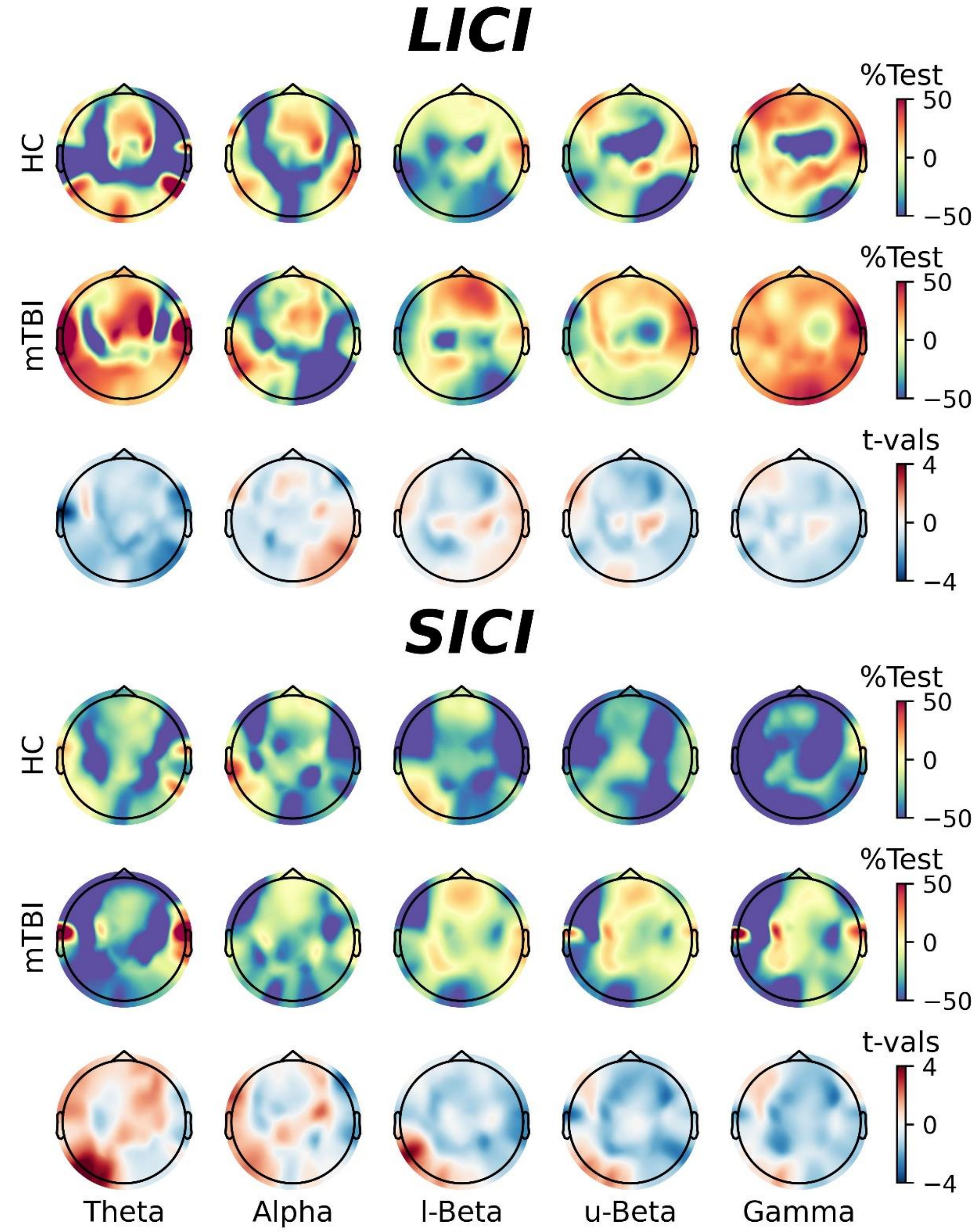


**Figure S5. Effects of mTBI on oscillatory indices of ICI without paired-pulse correction.** Normalised changes in TMS-related oscillations following application of LICI **(top panel)** and SICI **(bottom panel)** in healthy controls *(top row)* and mTBI patients *(middle row)*, in addition to results of cluster-based comparisons between groups *(bottom row)*. Data have not undergone the subtraction procedure to remove the TEP generated by the conditioning stimulus.
